# scTOP: physics-inspired order parameters for cellular identification and visualization

**DOI:** 10.1101/2023.01.25.525581

**Authors:** Maria Yampolskaya, Michael Herriges, Laertis Ikonomou, Darrell Kotton, Pankaj Mehta

## Abstract

Advances in single-cell RNA-sequencing (scRNA-seq) provide an unprecedented window into cellular identity. The increasing abundance of data requires new theoretical and computational frameworks for understanding cell fate determination, accurately classifying cell fates from expression data, and integrating knowledge from cell atlases. Here, we present single-cell Type Order Parameters (scTOP): a statistical-physics-inspired approach for constructing “order parameters” for cell fate given a reference basis of cell types. scTOP can quickly and accurately classify cells at a single-cell resolution, generate interpretable visualizations of developmental trajectories, and assess the fidelity of engineered cells. Importantly, scTOP does this without using feature selection, statistical fitting, or dimensional reduction (e.g., UMAP, PCA, etc.). We illustrate the power of scTOP utilizing a wide variety of human and mouse datasets (both *in vivo* and *in vitro*). By reanalyzing mouse lung alveolar development data, we characterize a transient perinatal hybrid alveolar type 1/alveolar type 2 (AT1/AT2) cell population that disappears by 15 days post-birth and show that it is transcriptionally distinct from previously identified adult AT2-to-AT1 transitional cell types. Visualizations of lineage tracing data on hematopoiesis using scTOP confirm that a single clone can give rise to as many as three distinct differentiated cell types. We also show how scTOP can quantitatively assess the transcriptional similarity between endogenous and transplanted cells in the context of murine pulmonary cell transplantation. Finally, we provide an easy-to-use Python implementation of scTOP. Our results suggest that physics-inspired order parameters can be an important tool for understanding development and characterizing engineered cells.

## I. INTRODUCTION

Humans and other mammals have many specialized cells that arise through a complex developmental process [1]. At the blastocyst/epiblast stages of embroynic development, cells are pluripotent and possess the flexibility to develop into fates from all three germ layers [2]. However, as embryos mature, cellular identity becomes less plastic and more specified through a complex spatiotemporal process involving signaling and other environmental cues. The questions of how and why cells end up in one fate or another are fundamental to developmental biology [3–5]. More practically, understanding the process of cell fate specification is crucial for developing directed differentiation protocols for engineering cells *in vitro*, with important clinical applications for human health and disease [6–8].

One powerful set of techniques for characterizing cellular phenotype is transcriptome-wide measurement using single-cell RNA sequencing (scRNA-seq). Typical scRNA-seq experiments involve thousands of genes and dozens to millions of cells. For this reason, this data is intrinsically high-dimensional and requires specialized computational tools for analysis and interpretation [4, 9, 10]. This task is especially difficult because the technical challenges of sequencing RNA from individual cells result in data that is noisy and sparse [11–13]. Genes with relatively low expression are often recorded as not being expressed at all, resulting in zero counts called dropouts. Experiment-to-experiment differences in sequencing methods, platforms, and other difficult-to-standardize conditions result in technical variations in scRNA-seq measurements called batch effects. These challenges merit careful consideration of how to normalize and analyze the data in a way that permits the study of biological variation without conflation by technical variation.

scRNA-seq is often used for cellular identification. The most frequently used workflow involves applying dimensional reduction (such as Uniform Manifold Approximation and Projection (UMAP) [14] or SPRING [15]) to visualize the data in two dimensions, then using unsupervised clustering to group individual cells. These groups are then often manually annotated according to expression levels of known marker genes. Machine learning classifiers such as support vector machines are also commonly used to identify cells after training with labeled data [16]. Another common analysis task is trajectory inference, where the goal is to infer developmental trajectories from static time shots (reviewed in [17, 18]). Trajectory inference arranges cells on a low-dimensional manifold and assigns pseudotimes according to transcriptomic similarity. There are also methods such as partition-based graph abstraction [19] that combine clustering and trajectory inference to simultaneously calculate pseudo-times and provide discrete labels at varying resolutions. RNA-velocity-based methods like Dynamo [20] use splicing information or time-resolved data to predict paths of differentiation from RNA data.

Despite the power and utility of these computational tools, most commonly employed methods for analyzing scRNA-seq data share certain limitations. Dimensional reduction is often one of the first steps in visualizing scRNA-seq data, but current methods highly distort cell-to-cell and cluster-to-cluster quantitative relationships [21]. Distorted embeddings inconsistently represent distances between individual data points and between cell types, making it difficult to interpret cell fate transitions. Similar issues arise in trajectory inference and RNA-velocity methods because local neighborhoods can also vary substantially with the choice of embedding. Another limitation of these methods is the lack of reproducibility inherent to stochastic machine learning methods like t-SNE and UMAP, which output different results on separate executions even when hyperparameters remain the same [22]. Many algorithms also perform feature selection (e.g., limiting their analysis to the most variable genes or highly-expressed genes) or require choosing parameters such as perplexity, making results even more sensitive to the inclusion or exclusion of different datasets in the analysis.

An ideal analysis algorithm for scRNA-seq data should possess certain desirable properties. First, the algorithm should incorporate explicit assumptions that can be kept consistent between analyses. Second, the algorithm should be deterministic, further enhancing reproducibility. Third, the axes in visualization should be interpretable, intuitive, and remain fixed across analysis and datasets. Fourth, the algorithm should assign a continuous value to cell identity instead of assigning discrete identity labels, thus enabling a more subtle and fine-grained analysis of cellular identity and developmental trajectories that reflect the continuous nature of biology. Finally, the algorithm should be easily able to incorporate datasets from multiple data sources, including the wealth of new single-cell atlases now being generated [23].

Here, we present a new physics-inspired algorithm, single-cell Type Order Parameters (scTOP), that has all of these desired properties. The algorithm is based on the statistical physics concept of an order parameter – a coarse-grained quantity that can distinguish between phases of matter. Arguably the most famous example of an order parameter is magnetization in a ferromagnetic system (i.e., a system that can spontaneously magnetize). In a ferromagnet, the magnetization measures the alignment of spins and can be used to distinguish magnetic and paramagnetic phases [24, 25]. It is straightforward to generalize the idea of magnetization to more complex settings such as spin glasses and attractor neural networks [26]. In the context of epigenetic landscape models of cell identity, this takes the form of generalized magnetizations that measure how aligned a gene expression state is with each of the attractor basins for different cell fates [27, 28]. These generalized magnetizations serve as natural order parameters for cellular identity and can be calculated directly from data [27–30]. The scTOP algorithm builds on previous work by developing methods for integrating scRNA-seq data, greatly expanding the power and utility of this approach.

The inputs to scTOP are the scRNA-seq measurements of query data as well as a reference basis containing the archetypal gene expression profiles for cell types of interest. The outputs of scTOP are the coordinates of the query data in cell type order parameter space (i.e., generalized magnetizations), with each output dimension serving as a measure of similarity between the query data and a distinct cell type in the reference basis. In this way, scTOP naturally projects gene expression deterministically onto a space with dimension equal to the number of cell types in the reference basis. These generalized coordinates can be used for both cell type identification and visualization. One appealing feature of scTOP is that the assumptions of the algorithm are explicit and are contained entirely in the choice of reference basis. This allows direct comparison across datasets and settings since the coordinates of and distances between data points do not change as long as one uses the same reference basis. One important limitation of scTOP worth noting is that it cannot be easily used to identify novel cell fates. The underlying reason for this is that scTOP assumes one already knows what cell types are of interest in the form of a reference basis and, for this reason, is less suited for exploratory work with undocumented cell types.

In this paper, we show that scTOP can be used to reliably assess cell identity and accurately classify cell type with high accuracy at the resolution of individual cells. We demonstrate this using mouse and human samples from lungs, brains, and other organs, including transplanted cells that were generated through the directed differentiation of pluripotent stem cells (PSCs). By analyzing lineage tracing data and embryonic development data, we also establish scTOP’s capacity to quantify transitions between cell types, including bifurcations between closely related cell fates. We also show that scTOP can be used to quantitatively assess transcriptional similarity of engineered and natural cells by analyzing murine pulmonary transplantation experiments.

We have implemented scTOP in an easy-to-use Python package, available on the Python Package Index and Github. The code written for the analyses in this paper is also available on Github here.

## II. METHODS

### A. Mathematical Details

To calculate cell type similarity scores, scTOP uses a linear algebra projection method inspired by the concept of cell types as attractors in a dynamical system [31]. This projection method has been applied previously to bulk RNA-seq samples, and we will summarize the theoretical background here [27]. A cell type is a state of the gene regulatory network. A natural way to represent attractors in this system is with an attractor neural network, where attractors correspond to different cell types. If we represent each gene as a node in a network with an associated value measuring gene expression (positive value representing high gene expression, negative value indicating low gene expression), then a cell type may be denoted by a vector of gene expression values *S*_*i*_, where *i* = 1, …, *G* spans the *G* genes being measured. The gene expression profiles corresponding to the *C* cell type attractors are also *G* dimensional vectors 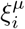, where *µ* = 1, …, *C* spans the *C* cell types being stored in the network. In what follows, we assume that *G > C*; in other words, the number of genes being measured is greater than the number of cell types.

Cell types are often highly similar in their patterns of gene regulation. Kanter and Sompolinsky [32] defined a pattern storage method for spin-glass-like neural networks where even correlated patterns robustly act as attractors. For the same system, they also defined order parameters: generalized magnetizations *a*^*µ*^, where *µ* iterates through each of the stored cell type attractors. The *a*^*µ*^ can be understood as de-correlated versions of the conventional spin glass magnetization order parameter that measures the similarity between a given network state and the network state *ξ*^*µ*^. Explicitly, the *C* generalized magnetizations *a*^*µ*^ can be calculated for a gene expression state *S*_*i*_ via the expression

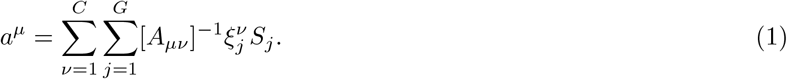

where

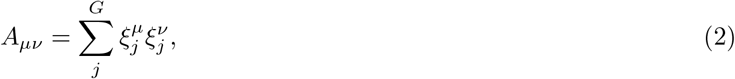

is the overlap matrix of gene expression profiles of different cell types. Previous works have shown the order parameters *a*^*µ*^ provided an excellent similarity score for analyzing bulk RNA-seq data [27–30]. However, bulk RNA-seq measurements average gene expression over many cells, rendering it impossible to measure cell fate transitions beyond an average tissue state.

scTOP uses the same order parameters, *a*^*µ*^, to track cell type at the resolution of individual cells, making it possible to directly observe cells in different stages of differentiation. In both attractor neural networks and previous applications to bulk RNA-seq data, the expression vectors *S*_*i*_ were binary variables. However, we have found that for working with scRNA-seq data, it is helpful to treat *S*_*i*_ as continuous.

### B. Preprocessing

As shown in figure 1 (a), the first stage of the algorithm is preprocessing and normalization of the input data. In our algorithm, measurements from each cell are normalized independently and do not depend on the expression profiles of any other cells in the dataset. We assume that the scRNA-seq data has been processed, resulting in a gene-wise count matrix: for each cell, each gene has an integer value that counts the number of assigned RNA reads. Genes are then rank-ordered. The rank order is subsequently converted to a z-score representing the percentile rank of the expression of a gene within the cell relative to all other genes being measured (assuming a normal distribution with mean zero and standard deviation one, for the sake of simplicity). For example, a gene that has higher expression than 97.8% of genes being measured is assigned a score of *z* = 2, while a gene that has higher expression than half of the genes is assigned a score of *z* = 0. The details of the process and its effects on data distributions are described in Appendix section IV A. The preprocessing step is a key component of the algorithm. By processing each cell independently, the output for one cell is independent of other cells included in the analysis. This is in contrast with other algorithms which normalize gene expression across cells by, for example, selecting the most variable genes.

**FIG. 1.**
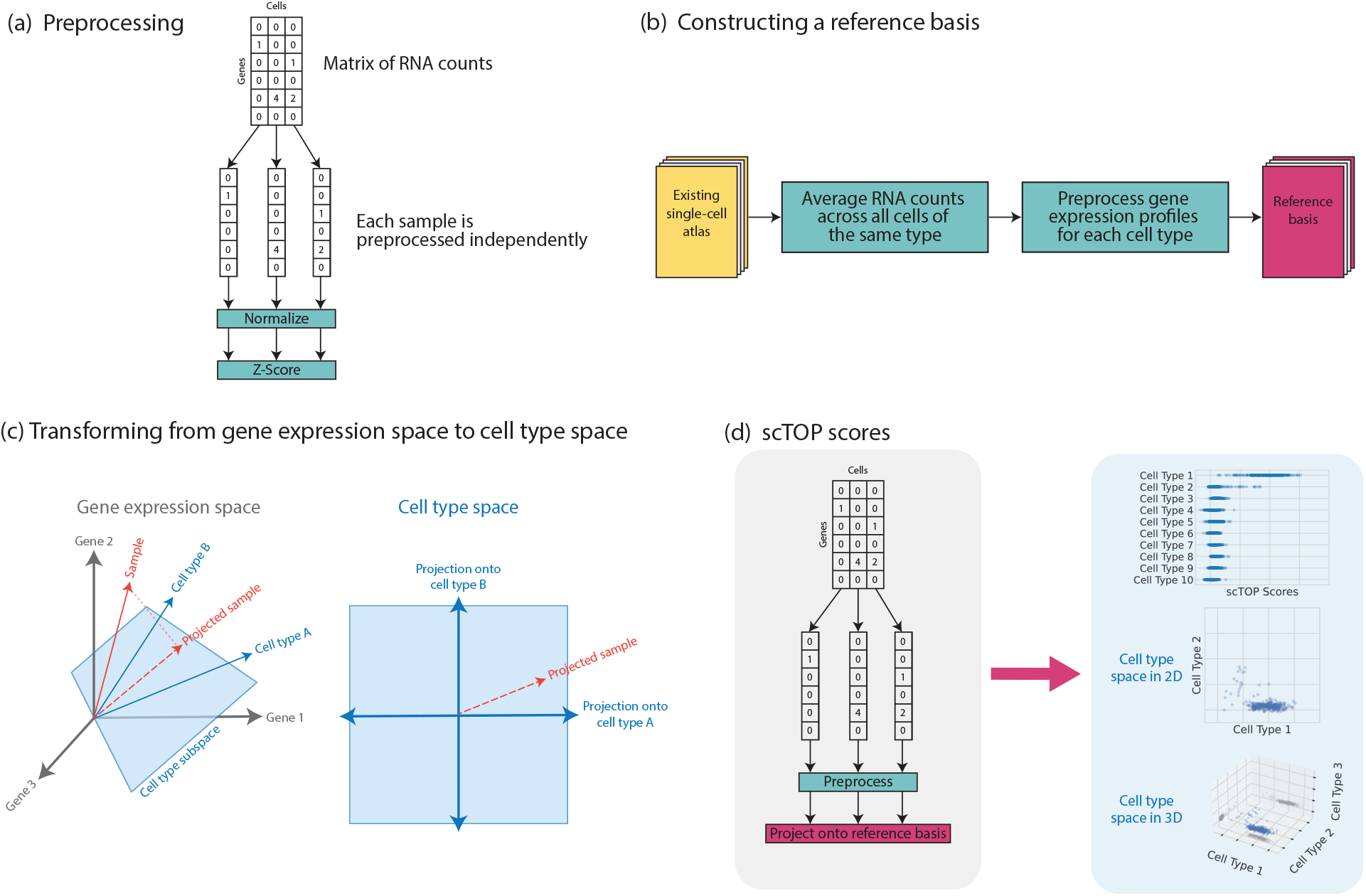
Steps involved in the scTOP algorithm. (a) Each cell in the query data is preprocessed independently of the other cells in the dataset. (b) scRNA-seq atlases, such as the Mouse Cell Atlas, are used to define the reference basis in the algorithm. (c)scTOP scores are the projections of query data onto the non-orthogonal subspace of cell types. Since there is no statistical fitting, no tuning parameters are involved. (d) The process of finding scTOP scores is shown in the grey section, and the blue section illustrates cell-type space. This space has as many dimensions as there are cell types in the reference basis. We can visualize this concept in 1, 2, or 3 dimensions, depending on which cell types are most relevant. In the 3-dimensional case, the shadows on the bounds of the scatter plot are included to better illustrate the position of points in 3-dimensional space.

### C. Construction of reference basis

Constructing a representative, accurate reference basis for the data being analyzed is vital to the scTOP algorithm. This process is shown in figure 1 (b). First, relevant single-cell RNA-seq atlases are gathered to be processed. For example, in analyzing mouse lung cells, we used the Mouse Cell Atlas [33], which contains single-cell samples of mouse tissues across all major organs. Once the relevant atlases are identified, we take an average across each cell population that corresponds to a distinct cell type. To mitigate the potent effects of noise and avoid training data imbalances, a sufficient number of cells must be included in each population. We have empirically found that the minimum sample size to achieve reasonable results is 100-200 cells. Noise affects the algorithm most strongly when it appears in the basis since the decorrelation operation involved in the projection separates cells according to the canonical expression levels included in the reference basis. After each cell population is averaged to create the gene expression profiles for the desired cell types, each cell type profile is preprocessed separately using the same procedure as that used for processing single cells: normalization followed by z-scoring. This creates a modular reference basis where cell types can easily be replaced, removed, or added at will.

### D. Calculation of scTOP scores

Given a reference basis, we can calculate the order parameters *a*^*µ*^ using Eq. 1. Although these order parameters were originally inspired by attractor neural networks and spin glass physics, the *a*^*µ*^ can also be understood as non-orthogonal projections onto the subspace spanned by the reference basis (i.e., cell type space). This is illustrated in figure 1 (c). A single-cell RNA-seq measurement of a sample can be thought of as a vector in *G*-dimensional gene expression space, where *G* is the number of genes. The *C*-cell types in the reference basis are also each represented by a *G*-dimensional vector and form a non-orthogonal basis for the *C*-dimensional cell type space (we assume *C < G*). To find the coordinates *a*^*µ*^ in cell type space, we project the sample vector onto the reference basis using Eq. 1.

Mathematically, we can always write the sample vector as a sum of the projected components and a component *S*_*i*_^⊥^perpendicular to the cell type subspace

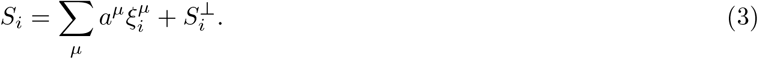

Biologically, *S*^⊥^ contains information about biological processes that introduce variation in transcriptional profile such as cell cycle as well as expression resulting from technical and biological noise. We refer to the *a*^*µ*^ as the scTOP scores. Each component of this vector measures the projection of the cell with gene expression profile *S*_*i*_ onto the *µ*-the cell type in the reference basis. These scores can be used to accurately classify cell identity and provide a natural visualization of gene expression in the space of possible cell fates (see figure 1 (e)).

In general, we are interested in calculating scTOP scores for individual cells. However, it is also sometimes useful to compute “aggregate” scTOP scores for cellular populations by averaging the mRNA counts *first*, then preprocessing the data. This produces a gene expression profile and a set of scTOP aggregate scores for each population. Aggregate scores tend to be higher because averaging over the RNA counts mitigates the well-documented effects of scRNA-seq noise [11, 12]; the averaged gene expression profile for the populations are not as sparse as individual profiles and thus are more similar to the averaged profiles that form the reference basis.

In the appendix, we give a detailed discussion of scTOP, including the effects of basis choice, score distributions for correct and incorrect cell types, and other technical details.

### E. Assumptions and computational complexity

scTOP does not have any stochasticity, require tuning of hyper-parameters, or statistical fitting procedures. The assumptions of the algorithm are explicit: that the gene expression profiles included in the reference basis are truly representative of the relevant cell types and that the reference cell types correspond to distinct cellular populations. We have found that constructing a reference basis for a cell type typically requires at least 100 cells. Additionally, if the cell types are not sufficiently different from one another, as in early embryonic development or certain stromal cells where the same cell type is compared between organs, the resulting scTOP scores may be unreliable and exhibit extreme sensitivity to small changes in the choice of reference basis. More information on the limitations of scTOP may be found in the appendix. Generally, outside these very limited edge cases, we have found that scTOP can reliably distinguish even extremely closely related cells (as shown in the example of pulmonary alveolar epithelial type 1 (AT1) and type 2 (AT2) cell lineages below).

Finally, we note that scTOP is extremely computationally efficient. The core of the scTOP is equation 1. This requires a single matrix inversion involving the reference basis that can be precomputed. For this reason, the computational complexity of scTOP scales *linearly* with the number of cells. The end result is that scTOP takes milliseconds to run, even for very large datasets.

## III. RESULTS

### A. Benchmarking and validation: Classification of cell fate

scTOP scores can be used to reliably and accurately predict cell identity. To show this, we applied the algorithm to several scRNA-seq datasets across species and laboratories. scTOP is organism-agnostic as long as an appropriate reference basis is used. Mice and humans are among the most common subjects of scRNA-seq measurements, and we restrict our analysis in this manuscript to data from these organisms. To do so, we make use of single-cell atlases, such as the Mouse Cell Atlas [33] and the Atlas of Lung Development [34].

Table I lists each of the datasets examined in this paper with their respective scTOP score accuracies. TopN refers to the percent of cells whose true cell types were in the top N scTOP scores, with a score greater than 0.1. Unspecified refers to the percent of cells for which the highest scTOP score was less than 0.1; this represents a failure of the algorithm to confidently identify the cell. The accuracies across species and tissues are high, and the unspecified percentages are very low, even in datasets exceeding a million cells.

**TABLE 1.** Accuracy scores of included sample types for each of the described data sets.

| Query Data | Species | Reference Basis | Top1 (percent) | Top3 (percent) | Unspecified (percent) | Total cells |
| --- | --- | --- | --- | --- | --- | --- |
| Kotton lab | Mouse | Mouse Cell Atlas | 97.19 | 99.77 | 2.69 | 4,805 |
| Brain atlas | Mouse | Self | 86.28 | 96.89 | 1.78 | 1,161,041 |
| CellBench | Human | Self | 100 | 100 | 0 | 2,822 |
| Lung atlas | Human | Self | 79.78 | 85.16 | 3.32 | 2,952 |
| Data set | Species | Reference Basis | Top1 (percent) | Top3 (percent) | Unspecified (percent) | Number of cell types |
| Mouse Cell Atlas | Mouse | Tabula Muris | 79.17 | 87.5 | 0 | 48 |

scTOP performs comparably to state-of-the-art methods for cell identification. Abdelaal et al. compared the performance of automatic cell identification algorithms for various datasets, including the CellBench dataset and an older version of the Allen Mouse Brain atlas [16]. They found that the median F1-score for the CellBench dataset ranged from 0.9 to 1; scTOP has a median F1-score of 1 for the same data. The version of the Allen Mouse Brain atlas they analyzed contained three levels of cell population annotations, with 3, 16, and 92 populations for each annotation level. The median F1-score ranged from 0.64 to 1 for the 16-population case and 0 to 0.98 for the 92-population case. The version of the brain atlas we analyzed contained 40 populations, and scTOP had a median F1-score of 0.88 for this dataset.

#### 1. Example 1: Lung lineages

To demonstrate the efficacy of scTOP in the case where the reference basis and query data come from different sources, we projected mouse lung data from Herriges et al. [35] onto the mouse cell atlas. The Herriges et al. dataset measures gene expression levels of six different specialized lung types from healthy mice: AT1, AT2, basal, ciliated, secretory, and neuroendocrine (See figure 2). The reference basis used to find the individual scTOP scores for this data set was derived from the Mouse Cell Atlas. Even though the Herriges et al. dataset and the Mouse Cell Atlas were created in different laboratories using different sequencing methods, the resulting scTOP scores distinctly separate the Herriges et al. dataset into the expected cell types. Furthermore, the individual scTOP scores strongly correlate with the expression of lineage marker sets (figure 2). Altogether this suggests that scTOP provides a reliable method to annotate scRNA-seq datasets without the need for the expertise required to generate lineage marker gene sets.

**FIG. 2.**
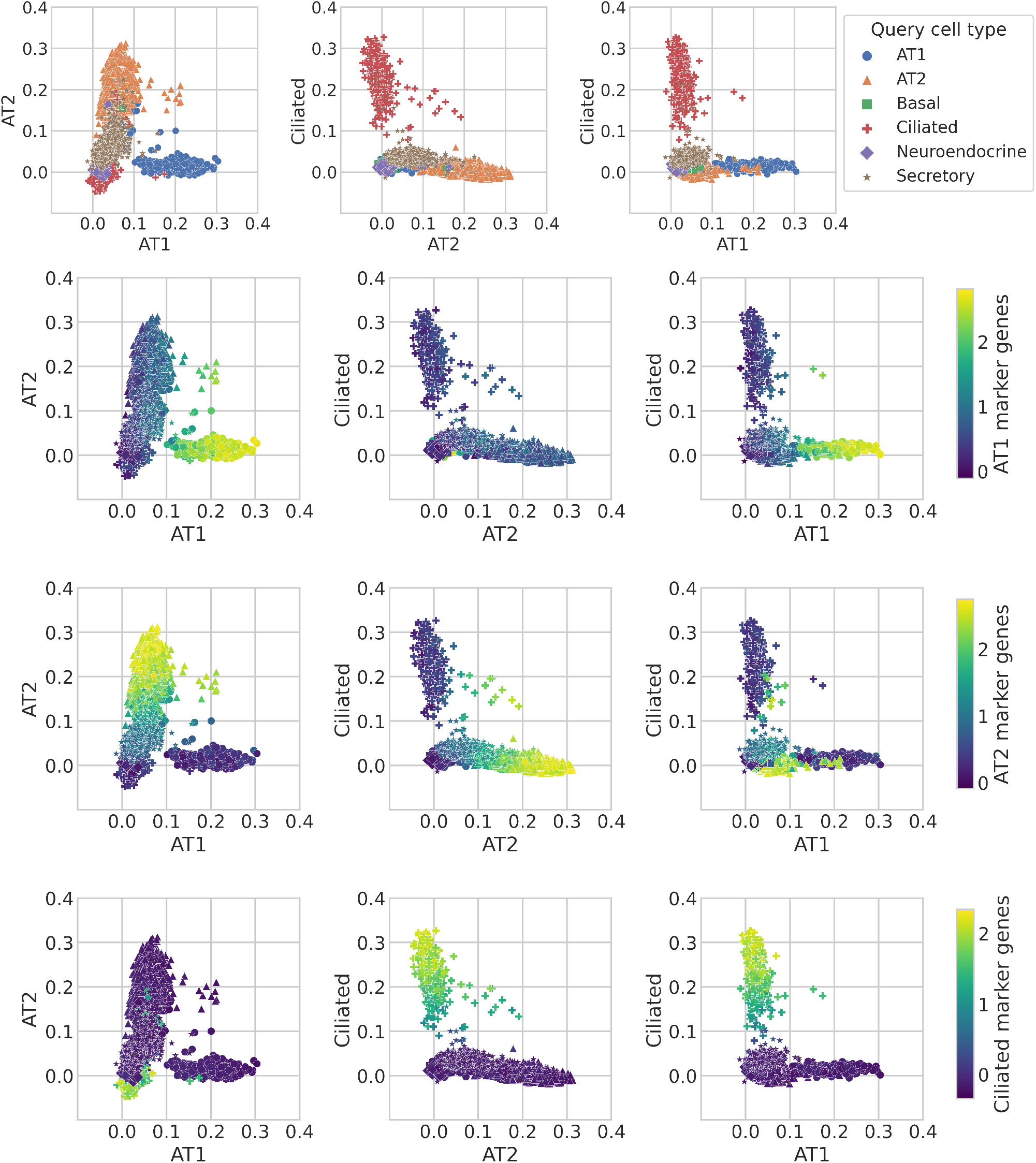
scTOP identifies mouse lung cell types. Using the Mouse Cell Atlas as the reference basis, we show that mouse lung types separate clearly on the scTOP score axes. The data points correspond to individual cells, and in the first row of plots, the marker shape and color indicate the true cell type as determined by annotations from Herriges et al. The axes used are scTOP scores for similarity with lung ciliated cells and alveolar types 1 and 2. The second, third, and fourth rows of plots have individual cells colored by the marker genes for the type indicated by the color bars at the end of each row (see appendix IV E for details on marker genes used).

#### 2. Example 2: Mouse Brain Atlas

The mouse brain atlas [36] sequenced over one million cells and clustered them into 42 subclasses and 101 supertypes. For a reference basis, we reserved a subset of 200 cells from each of the identified subclasses to use as training data. The test data, from which the accuracy score was calculated, was the remaining cells of each type not included in the training data. Ten different cell types are visualized in a grid in figure 3 (a). The color of each grid square corresponds to the average scTOP score of cells annotated as the type on the y-axis, with the cell type corresponding to the scTOP dimension on the x-axis. As shown by the distinct blue diagonal, the average scTOP score correctly matches the corresponding annotated cell type. This demonstrates that scTOP is able to distinguish cell fates even from closely related neural lineages.

**FIG. 3.**
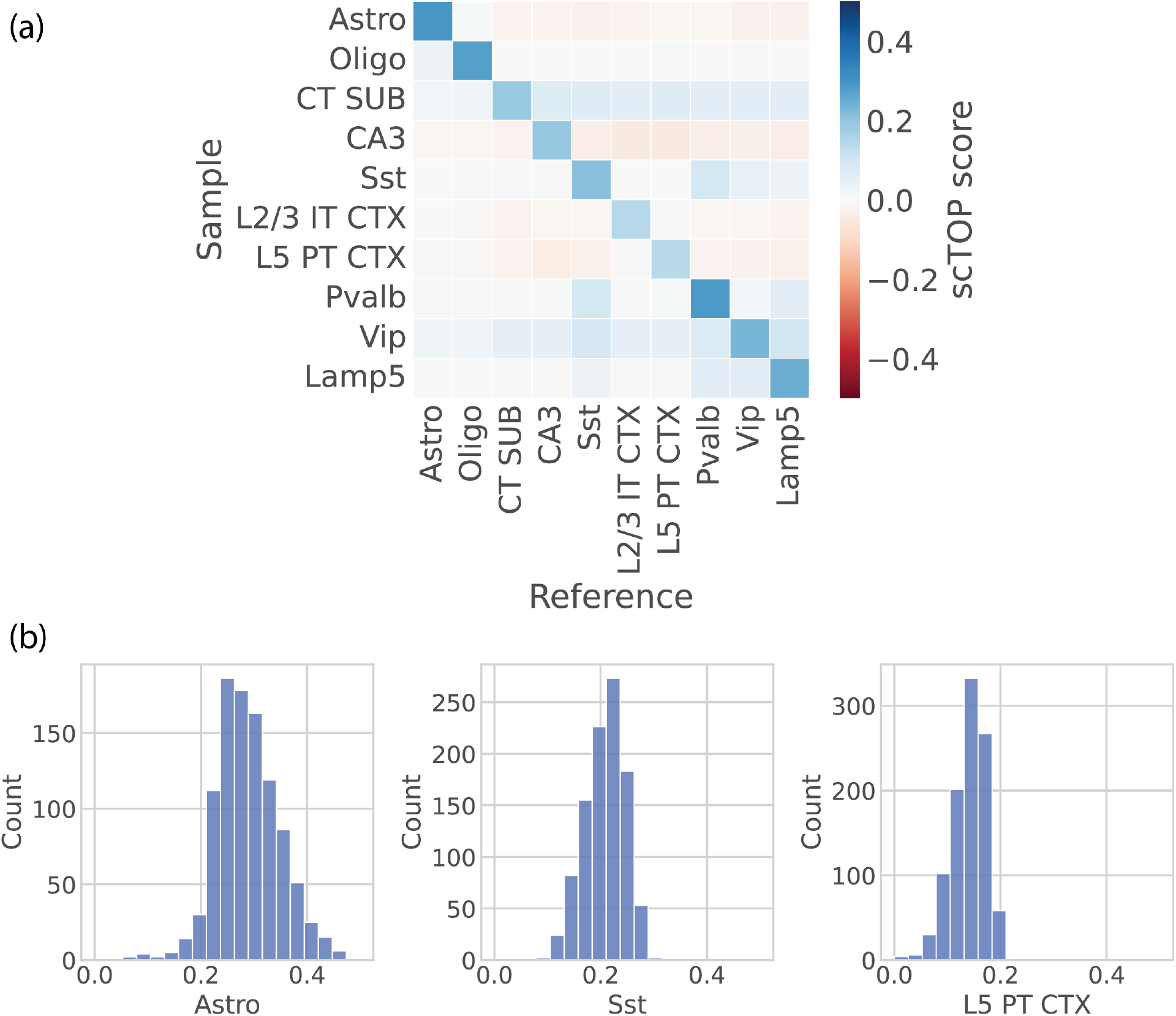
scTOP identifies mouse brain cell types. (a) Heatmap showing the average scTOP score for individual cells of the type indicated on the y-axis, compared to the reference types on the x-axis. A subset of cells from the Mouse Brain Atlas is used as the reference basis to query other cells from the same data set. The diagonal indicates that scTOP accurately matches query cells to the true reference type. (b) Histograms showing the distribution of scTOP scores for individual cells of the type labeling the x-axis. The scores shown are of the same type as the sample data. The far left histogram shows the distribution of Astro scores for Astro cells, and the average of this distribution is indicated by the color of the upper left corner box in the heatmap shown in (a).

#### 3. Example 3: Matching cell types to tissues

scTOP can also be used to match cells with tissues/organs. We illustrate this by comparing cells from the Mouse Cell Atlas with a reference basis consisting of Tabula Muris organs [37]. To do so, we averaged RNA counts across cell populations from the Mouse Cell Atlas, then preprocessed the averaged data and calculated scTOP scores. These aggregate pseudo-bulk scores tend to be higher than individual cell scores because averaging over many cells compensates for the dropout effect. As shown in figure 4, we found the aggregate scTOP scores for cells from the Mouse Cell Atlas were correctly and strongly identified with the annotated tissue of origin in the Tabula Muris atlas. Parenchymal cell types were the most associated with the correct organ, while stromal cells were sometimes misidentified – for example, the mammary gland endothelial cells. This suggests that stromal cells are less specified to each organ. A discussion of the accuracy of scTOP, when applied to stromal types versus parenchymal types, may be found in appendix IV D 1.

**FIG. 4.**
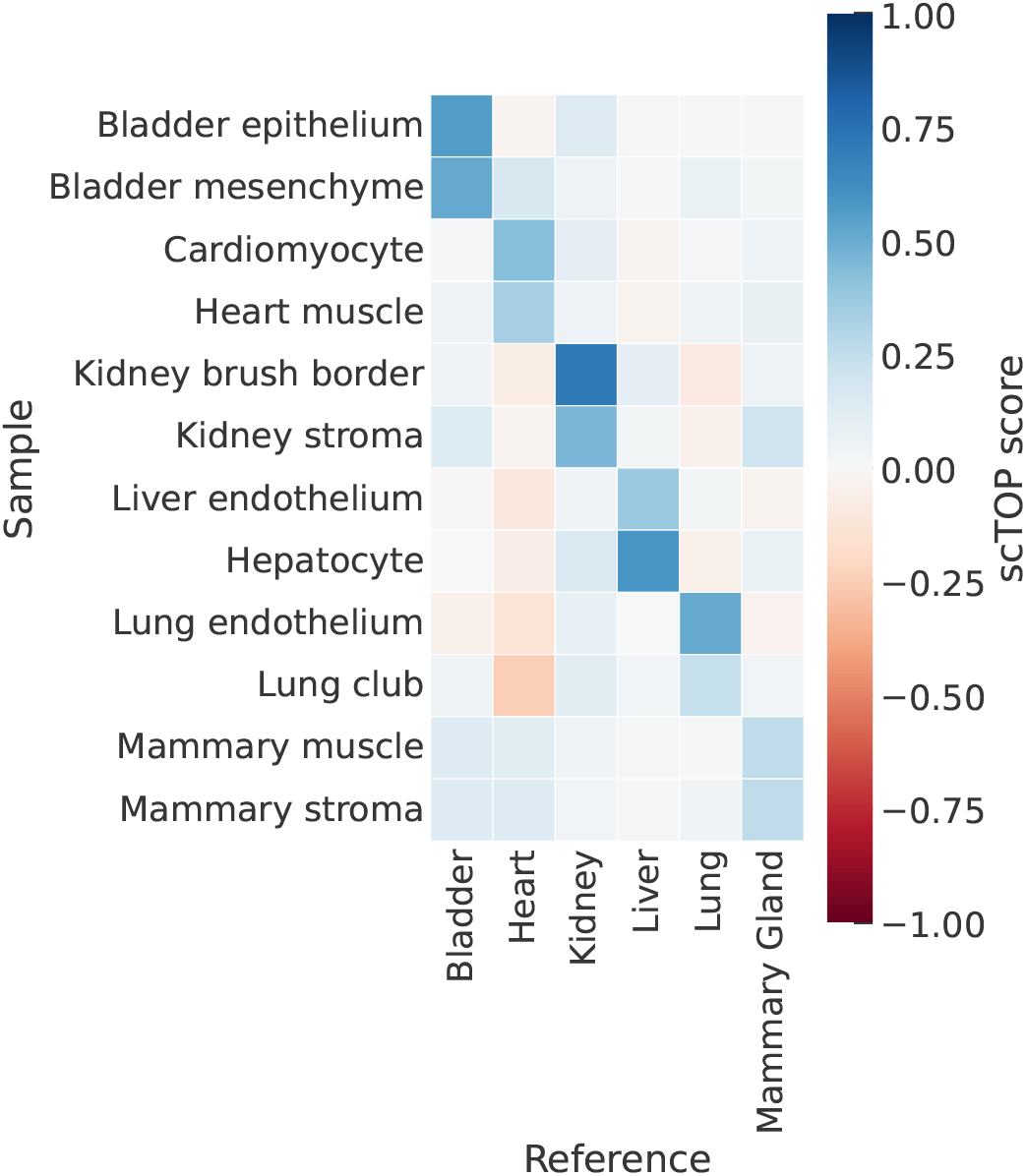
scTOP is able to classify tissues at different resolutions. Aggregate pseudo-bulk samples from the Mouse Cell Atlas are compared to a reference basis consisting of organs from Tabula Muris. The scTOP scores are significantly higher when comparing a cell type with the organ of origin. The shade of each bin indicates the average scTOP score for an individual queried cell compared to the cell type indicated on the x-axis. The y-axis lists the true types of the queried cells. The dark diagonal demonstrates the scTOP scores correctly match the queried data to the corresponding true types.

#### 4. Example 4: Human data

The CellBench dataset [38] was designed specifically to benchmark scRNA-seq analysis methods. We used the subset of data pertaining to five cancer lines. 200 cells from each line were used as training data to create the reference basis, then the rest of the cells were input as query data. As shown in figure 5 and table I, scTOP classifies the cell line identity of the test data with 100 percent accuracy.

**FIG. 5.**
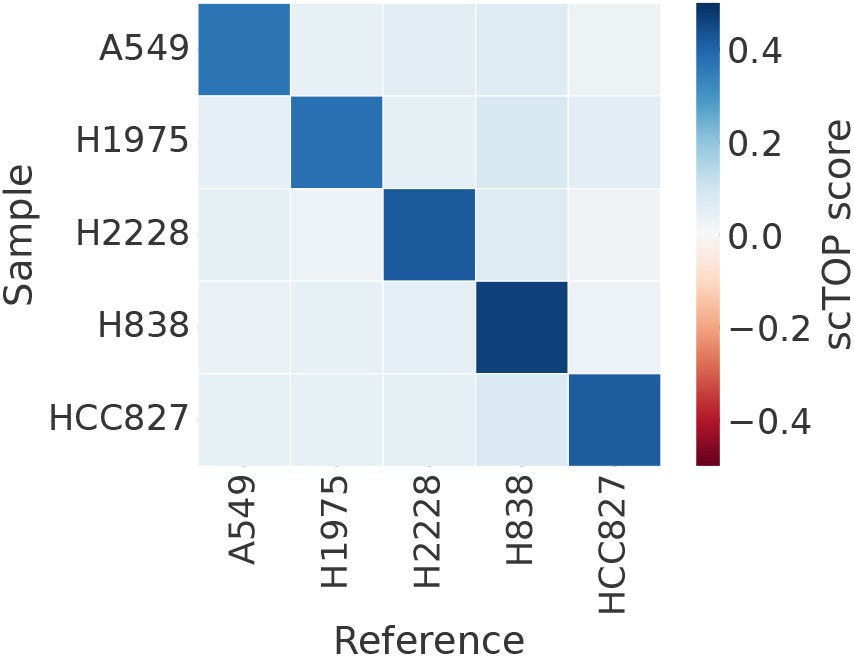
scTOP correctly identifies human tissues. The CellBench data set contains human cell cancer lines specifically processed to benchmark scRNA-seq algorithms. We take a subset of cells to use as the reference basis in analyzing the rest of the CellBench data. The shade of each bin indicates the average scTOP score for an individual queried cell compared to the cell type indicated on the x-axis. The y-axis lists the true types of the queried cells. The dark diagonal demonstrates the scTOP scores correctly match the queried data to the corresponding true types.

To illustrate the power of scTOP for classifying human samples, we analyzed data from the human lung atlas dataset [39]. This dataset sequenced thousands of human lung cells and identified 58 phenotypic populations. For our analysis, we restricted the training and test data to epithelial and stromal types that had at least 200 cells in each type population. The reference basis was created using 80% of the cells from each population, and the query data consisted of the remaining 20% of the data. Figure 6 shows that the scTOP scores were consistently high in cases where the score type matched the true type of the query.

**FIG. 6.**
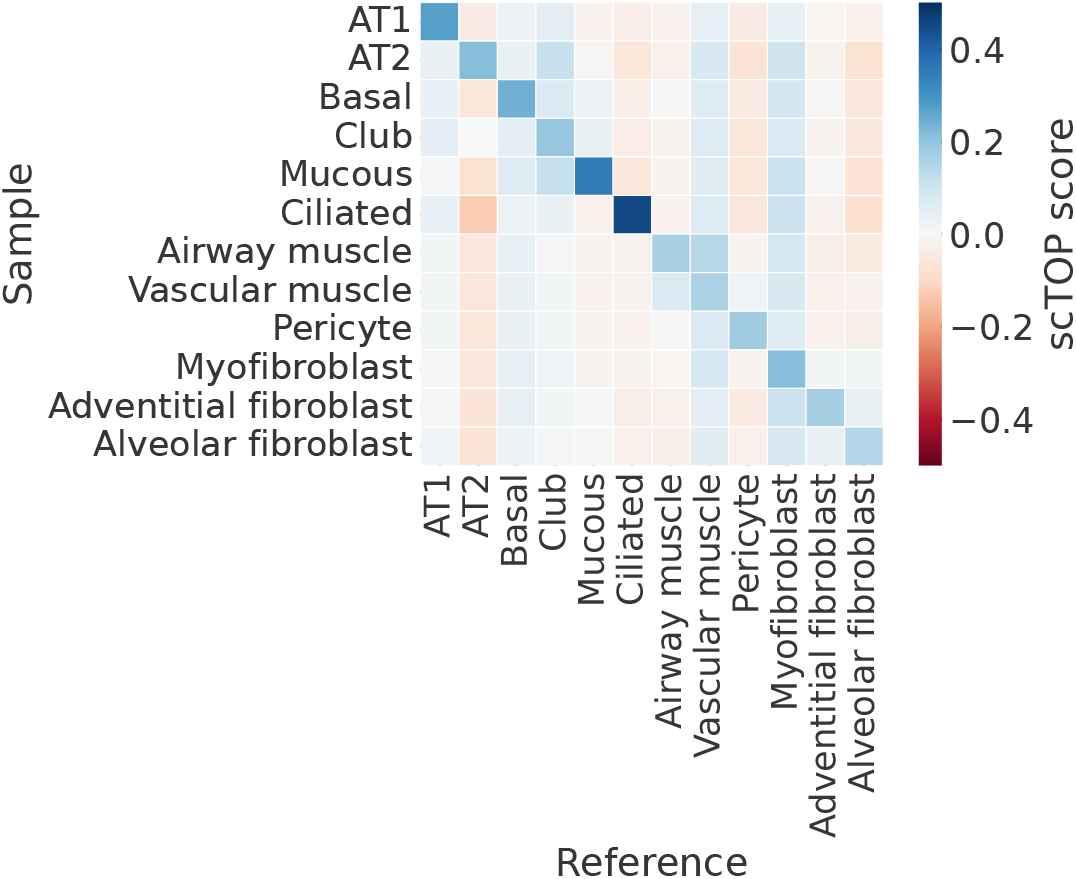
scTOP correctly identifies human tissues. Similar to figure 5, cells from the Human Lung Atlas are compared to a reference basis constructed from cells from the Human Lung Atlas. The colors of the bins correspond to the average scTOP scores of individual cells. Again, the diagonal is the most prominent section, showing that cells of the y-axis types are correctly matched to the reference x-axis identities.

As demonstrated by the diverse data sources examined here, as long as a well-defined reference basis is available, scTOP works extremely well for cell type identification across different measurement conditions, species, organs, and resolutions. This performance is especially impressive since, as discussed earlier, the algorithm does not use any statistical fitting procedures or dimensional reduction methods such as PCA, tSNE, UMAP, or SPRING.

### B. Visualizing developmental dynamics using scTOP

Having demonstrated the robust ability of scTOP to identify stable cell types, we next demonstrate how scTOP can be used in combination with scRNA-seq time series to visualize transient cell states in dynamic processes like development and differentiation.

#### 1. Lung development

The alveolar epithelium of the lung undergoes dramatic maturation and morphological changes during embryonic and perinatal development, ultimately giving rise to AT1 cells and AT2 cells. To investigate this process with scTOP, we reanalyzed alveolar epithelial cells from a recent dataset by Zepp et al. [40], which includes seven time points between embryonic day 12.5 (E12.5) and postnatal day 42 (P42). These cells were then compared against a reference basis developed from adult cell types in the Mouse Cell Atlas and an E12.5 epithelial progenitor cell type from the Atlas of Lung Development [34].

The resulting AT1 and AT2 scTOP scores were plotted to visualize maturation into these two lineages (Figure 7(a)). As expected, early embryonic progenitor cells (E12.5, E15.5), which express few of the mature lineage markers, align poorly with both lineages. In contrast, the E17.5 alveolar epithelial cells fall along a wide spectrum with cells aligning to AT1 cells, AT2 cells, or both lineages. In the postnatal time points, this continuous spectrum gives way to more distinct clusters, eventually resolving into AT2 and AT1 cell clusters at P42. Similar to our analysis of adult mouse lungs (Figure 2), the scTOP scores seen over development correlate well with our lineage marker gene sets (Figure 7 (b)-(c)).

**FIG. 7.**
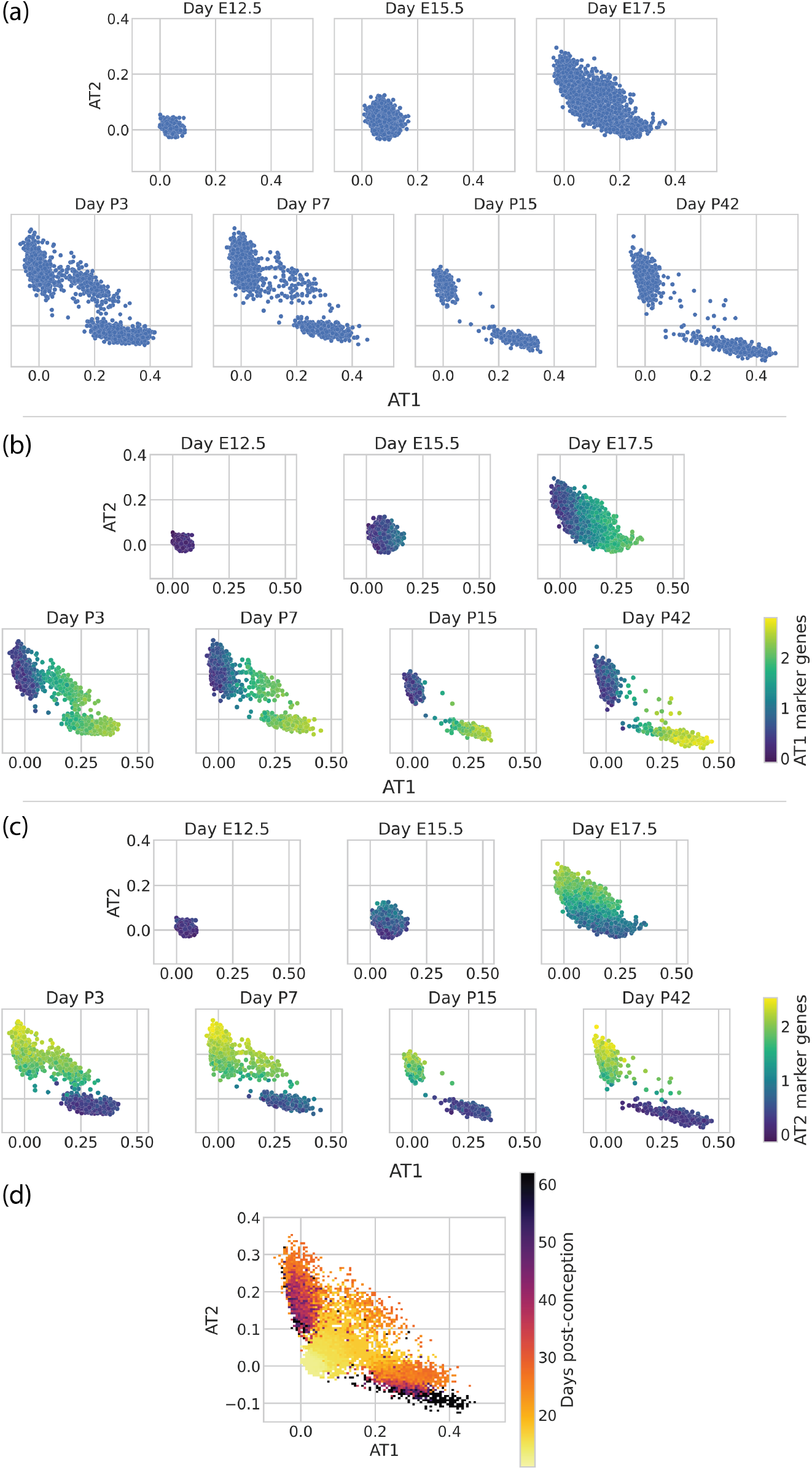
Visualizing the developing mouse lung with scTOP. scTOP illustrates the specification of alveolar type I and type II cells in murine embryonic development. Alveolar cells from Zepp et al. are compared to a reference basis constructed from adult lung cells from Herriges et al. and early epithelial cells from the Atlas of Lung Development. (a) Each subplot corresponds to the age of the mouse from which the sample was extracted, from post-conception day 12,5 (E12.5) to 42 days post-birth (P42). (b), (c) Scatter plots of the same data as in (a), now with each cell colored by the average over that cell’s expression of (b) AT1 marker genes and (c) AT2 marker genes. The scTOP scores of AT1 (AT2) cells are high when the average AT1 (AT2) marker gene expression is high. (d) The same data as above is displayed as a 2D histogram, with each bin colored corresponding to the average day of the data point(s) falling within that bin. The color trend shows that early cells generally have very low AT1 and AT2 scores, then the AT1 and AT2 scores increase over time.

In addition to the expected postnatal AT2 and AT1 cell clusters, this scTOP analysis illustrated a transient AT2/AT1 hybrid population that persisted through P7. The existence of these hybrid AT2/AT1 cells was already noticed in Zepp et al. using a combination of graph-based clustering and gene enrichment profiling (see Figure 1C, I in [40]). The authors conjectured that these AT2/AT1 cells were a transitional state of AT2 similar to Spock2+/Axin2+ AT2 cells previously identified in adult lungs [41]. To assess whether this was the case, we displayed Axin2 expression on our scTOP plots (Figure 8(a)). These cells do not have significant upregulation of Axin2 relative to perinatal or adult AT2 cells, suggesting that the postnatal hybrid AT2/AT1 cells are distinct from adult Spock2+/Axin2+ AT2 cells [41].

**FIG. 8.**
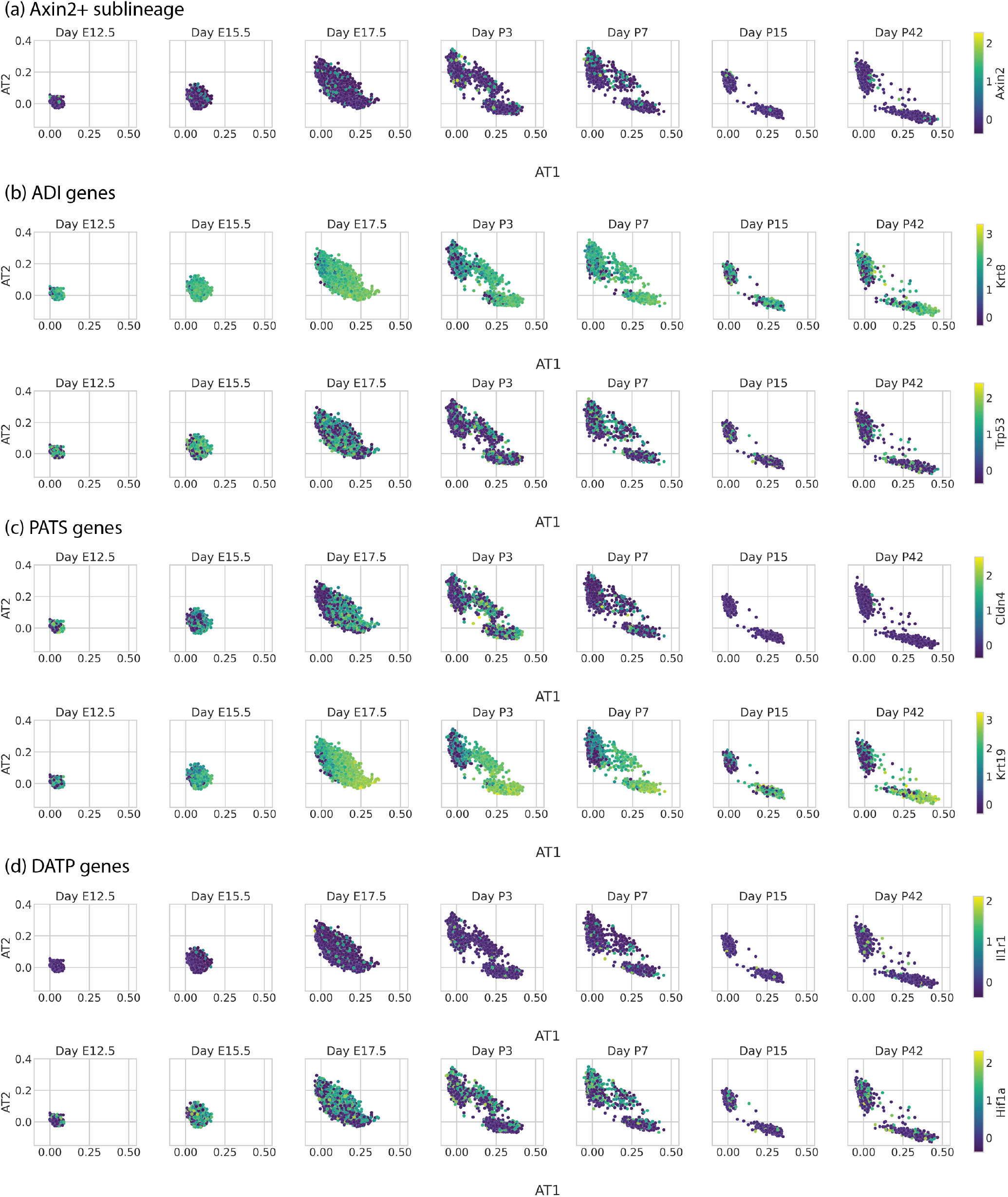
The AT1/AT2 combination state (most apparent in days P3 and P7) do not uniquely express genes of AT1/AT2 transitional states described by previous papers. Alveolar cells from Zepp et al. are plotted on AT1 and AT2 axes, separated by day. Each cell is colored by the expression z-score of a particular gene, and the particular gene is different in each row. The gene used to color the cells is indicated by the color bar on the far right of each row. (a) Alveolar cells colored by Axin2, showing that the AT1/AT2 combination cells do not appear to belong to the Axin2+ sublineage described by Frank et al. [41]. (b) Alveolar cells colored by Krt8 and Trp53 expression, which are genes corresponding to alveolar differentiation intermediate cells (ADI) described by Strunz et al. [43]. AT1, AT2, and the combination AT1/AT2 state all express Krt8 and Trp53 at similar levels. (c) Alveolar cells colored by expression of pre-alveolar type-1 transitional cell state (PATS) genes Kobayashi et al. [44]. AT1, AT2, and the combination AT1/AT2 state all express Cldn4 at similar levels. AT1 and AT1/AT2 combination cells both express Krt19 at similar levels, although AT1 cells express the gene at higher levels. (d) Alveolar cells colored by expression of damage-associated transient progenitor (DATP) genes described by Choi et al. [45]. AT1, AT2, and the combination AT1/AT2 state all express Il1r1 and HIF1a at similar levels.

We then performed a similar analysis of markers for other published postnatal hybrid AT1/AT2 or transitional alveolar cell states, including alveolar differentiation intermediate (ADI, *Krt8* and *Trp53*), pre-alveolar type-1 transitional cell state (PATS, *Cldn4* and *Krt19*), and damage-associated transient progenitors (DATPs, *Ilr1* and *Hif1a*) [42–45]. In each case, markers were not uniquely upregulated in the perinatal hybrid AT1/AT2 cells relative to either the perinatal or adult committed lineages (Figure 8 (b) - (d)). Furthermore, differential gene expression analysis failed to identify any genes that were uniquely differentially expressed in the hybrid AT1/AT2 cells relative to either timepoint-matched AT1 or AT2 cells. These results suggest that this population is most likely transcriptionally distinct from previously identified adult AT2-transitional cell types.

#### 2. scTOP for hematopoiesis using lineage tracing data

In the previous section, we used scTOP to visualize differentiation into two closely related lung epithelial cell fates. In this section, we show how scTOP can also be used to gain insights into more complicated developmental processes such as hematopoiesis, in which hematopoietic stem cells (HSCs) give rise to multiple differentiated blood cell types. To do so, we make use of the unique dataset by Weinreb et al. [46] on hematopoiesis in mice using a new technique, lineage and RNA recovery (LARRY), that combines scRNA-seq and lineage tracing data. LARRY uses heritable barcodes that can be detected using scRNA sequencing, allowing for the tracking of clone families during differentiation. Here, we focus on visualizing the ex-vivo experiments in [46], where cells were sequenced at a sufficient depth to infer lineage relationships. In these experiments, HSCs were extracted from mice, barcoded, and allowed to expand in primary culture. scRNA-seq measurements were performed on days 2, 4, and 6 post-barcoding. The cells were then annotated using marker genes into 9 differentiated blood cell fates.

We restrict our analysis to a subset of these annotated cell types with at least 150 cells. We also excluded cells labeled as erythroids from our analysis since we found that this population was highly heterogeneous and gave rise to poor projections (likely due to the fact it was annotated using a single marker gene, *Hbb-bs* [46], which did not sufficiently define a distinct population). Furthermore, we treated all day 2 undifferentiated cells as a single population we called progenitors. This resulted in a reference basis consisting of seven cell types: undifferentiated progenitors, neutrophils, monocytes, megakaryocytes, mast cells, eosinophils, and basophils. To construct the basis, we averaged the RNA counts for 150 random cells of each of these types, only including cells if they had no sister clones. Then we preprocessed the averaged gene expression profiles and inserted them into the reference basis.

Figure 9 shows the differentiation dynamics of three different clonal families using scTOP. The histograms show the distribution of scTOP projection scores on each cell type for days 2, 4, and 6 post-barcoding, while the scatter plots show two-dimensional cross-sections of scTOP projection scores as a function of time for the indicated cell types. In all the plots, the scTOP projections for the undifferentiated hematopoietic progenitor stem cell decreased over time, with a high projection on day 2 and a low projection on day 6, while the scores for the mature types increased in time.

**FIG. 9.**
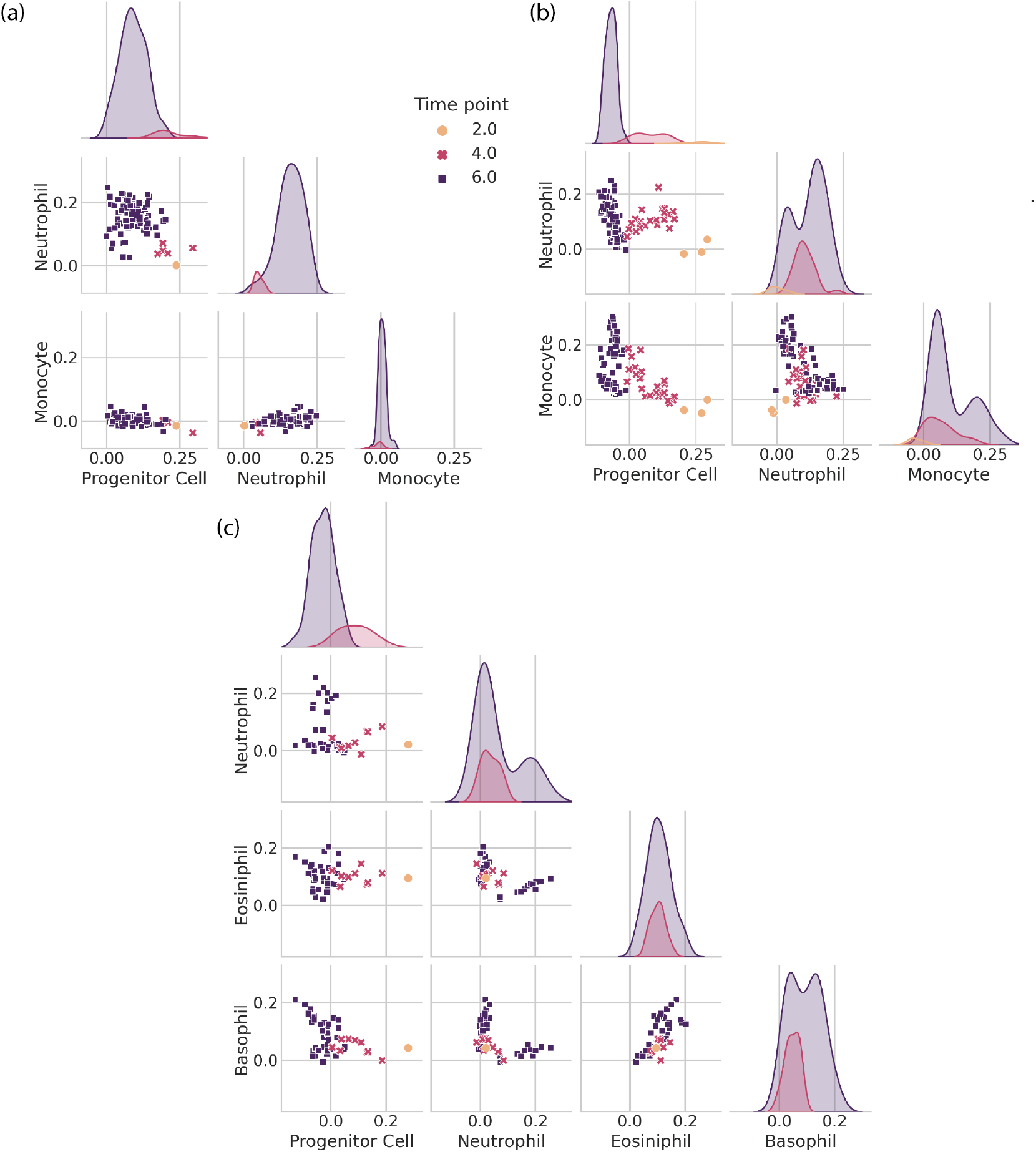
Tracing hematopoietic cell differentiation with scTOP. (a), (b), (c) display scatter plots and distributions of scTOP scores for individual in vitro clone families from [46]. The scatter plots show the x-axis scTOP score compared to the y-axis scTOP score for individual cells, while the distributions show the scTOP scores of the type indicated on the x-axis of the corresponding column. The color and shape of each marker (and the color of each distribution) indicate the days in culture on which the cells were extracted. (a) A clone family where all of the cells end up becoming neutrophils. (b) A clone family where some clones end up becoming neutrophils and some become monocytes. At day 2, the 3 sister cells have low neutrophil and monocyte scores and high progenitor scores. At days 4 and 6, the progenitor scores decrease, and a portion of the cells increase in monocyte scores while others increase in neutrophil scores. (c) A clone family where the descendant cells become neutrophils, eosinophils, and basophils.

Interestingly, the three clones show very different behaviors as they mature. Figure 9 (a) shows a clone where a single barcode detected at day 2 gives rise to multiple cells in the neutrophil lineage. This can be seen in the plots by noting that the day 6 cells have near zero scores on both progenitor cells and other differentiated lineages, such as monocytes, but project significantly on the neutrophils. Figure 9 (b) shows a barcode that was detected in three cells on day 2 that gives rise to a mix of neutrophils and monocytes on day 6. Interestingly, the day 4 cells move in the direction of both of these lineages before bifurcating at day 6. This data is consistent with Weinreb et al. [46], who previously highlighted the existence of such a bipotent neutrophil-monocyte progenitor population. We also find evidence that progenitor cells can even give rise to three distinct cellular populations. Figure 9 (c) shows a single barcode found on day 2 that results in day 6 cells with three distinct populations with non-zero scores on basophils, eosinophils, and neutrophils, suggesting a single clone can bifurcate into three different types of differentiated cell types.

Figure 9 focuses on scTOP scores for individual clonal families, while Figure 10 shows the scTOP projection scores across all 1,816 multi-generational clonal families in the dataset. In each subplot, the horizontal axes show progenitor scores, and the vertical axes correspond to the six differentiated cell types included in the reference basis. Each bin of the 2-dimensional histogram is colored according to the *average* day of all the cells that fall within that bin. For this reason, although the cells were only sampled on days 2, 4, and 6, the color bar is continuous, with a range between 2 and 6 days. The histograms show that, as expected, the day 2 cells generally have high projections on progenitor cells, but as time passes, progenitor scores decrease while the scores on differentiated cell types either increase (indicating cells that differentiate into the cell type on the y-axis) or remain close to zero (indicating cells that differentiate into a different cell type than that on the y-axis). Somewhat surprisingly, the plots for different cell types look qualitatively different from each other (for example, compare the plots for megakaryocytes and monocytes).

**FIG. 10.**
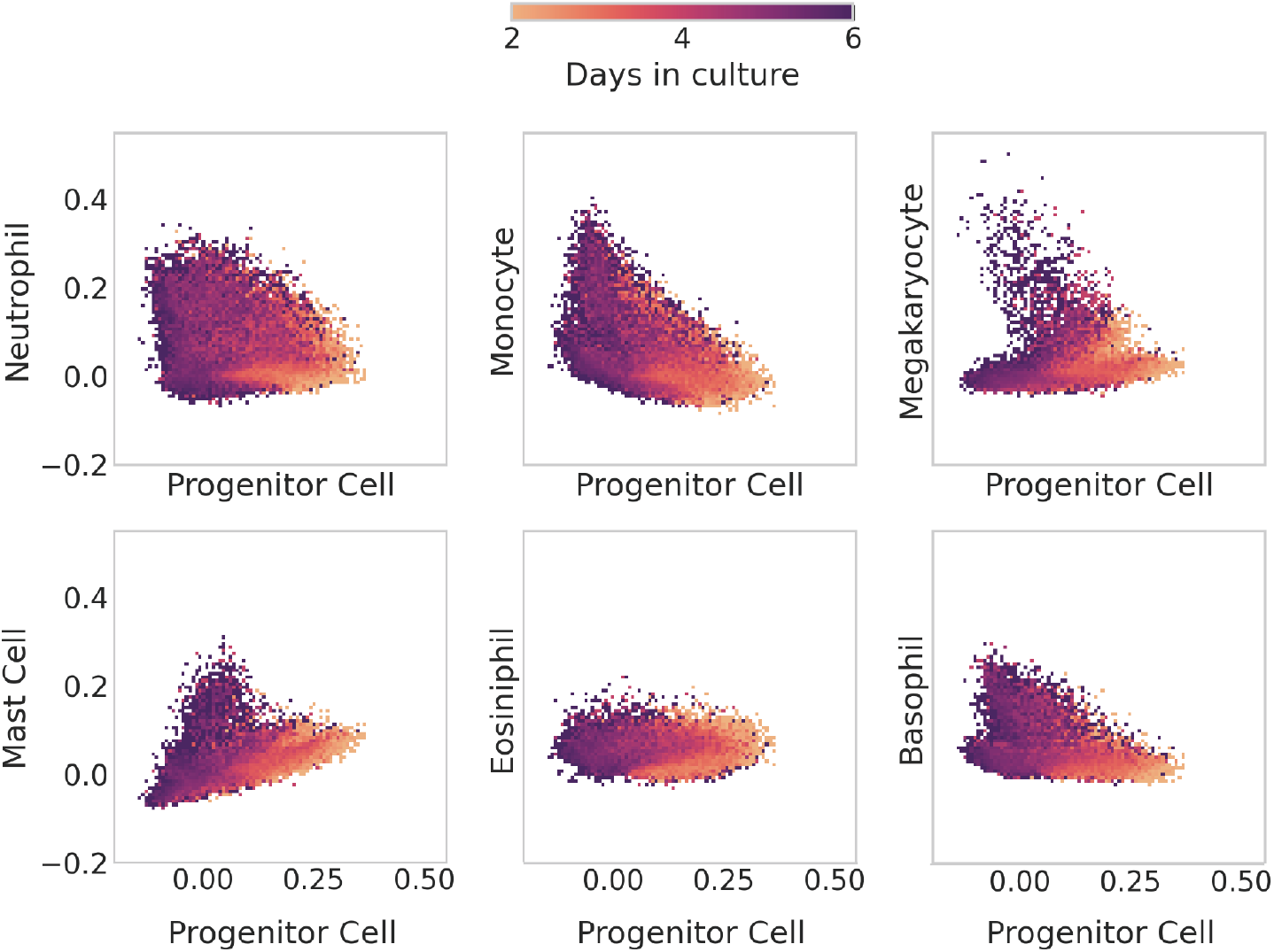
Tracing differentiation of hematopoietic cells from progenitor to mature type. The plots shown here are 2D histograms of all of the in vitro clone families from Weinreb et al., with the x-axis of each one indicating the scTOP score for hematopoietic progenitor type and the y-axis showing the score for a mature type. Each bin of the histogram is colored by the average day in culture of the data points that fall within the bin. Early cells have high progenitor scores and low mature-type scores. As time passes, cells end up with high scores in mature types.

To try to better understand this, we made additional 2-dimensional histograms comparing scTOP scores between all differentiated cell types in our basis (see Figure 11). What is striking about these plots is that they show two qualitatively different behaviors depending on which pairs of cells are being compared: the differentiation pathways for some pairs are mutually exclusive, while other pairs show a more graded behavior. For example, the histogram for megakaryocytes and monocytes exhibits a distinct L-shape, indicating that these developmental programs are mutually exclusive since cells never have significant projection scores of both cell types simultaneously. This is in stark contrast with the histogram for monocytes and neutrophils, where there exists a continuous spectrum of cells that are monocyte-like, neutrophil-like, and every step in between.

**FIG. 11.**
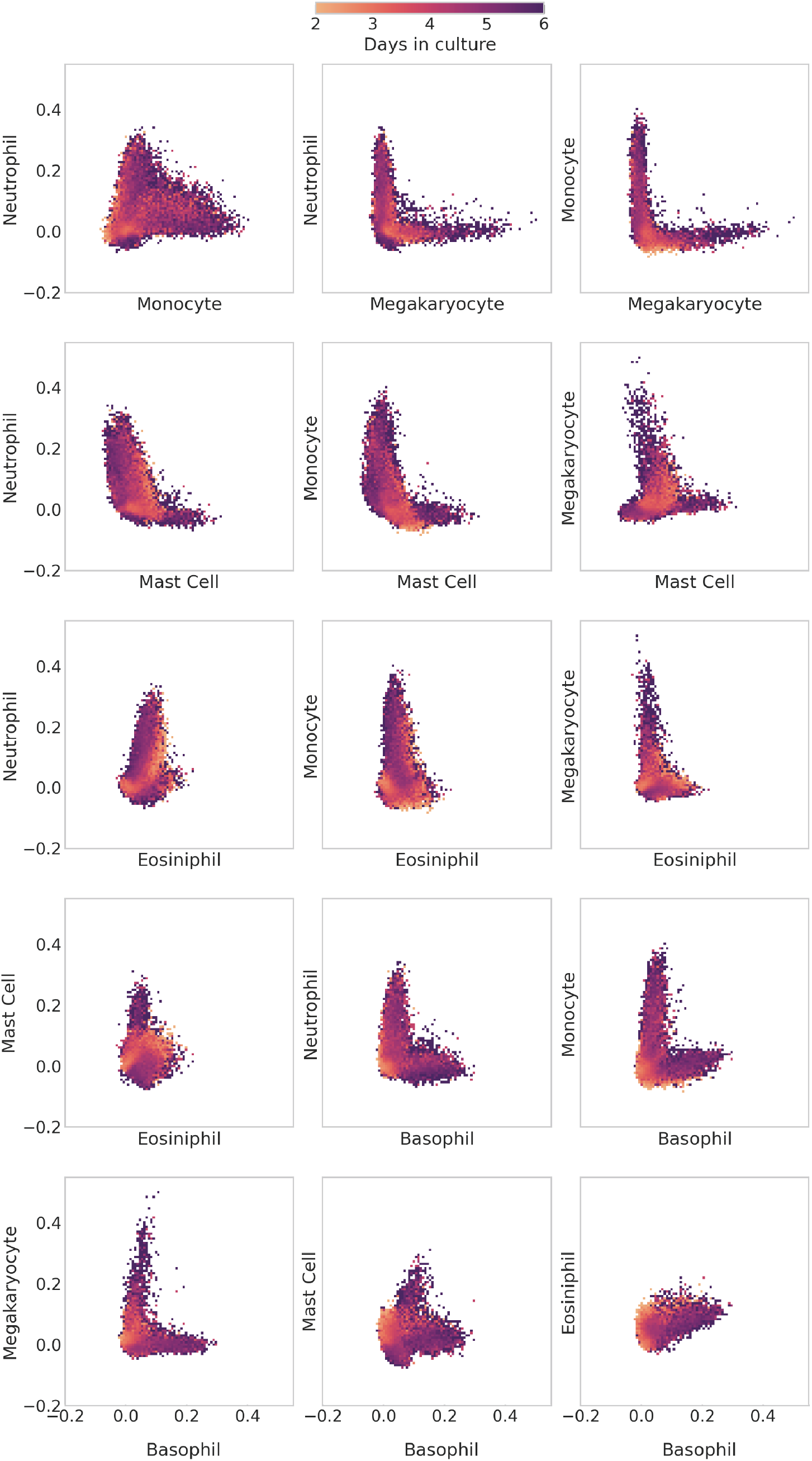
Comparing differentiation of hematopoietic cells between mature types. As in figure 10, the plots shown here are 2D histograms of all of the in vitro clone families from Weinreb et al. The x-axis and y-axis of each histogram indicate the scTOP score of the labeled mature type. Each bin of the histogram is colored by the average day in culture of the data points that fall within the bin. Some pairs of types have cells with high scores of both types. This is apparent in the neutrophil-monocyte histogram, which presents a continuous spectrum of scores from the horizontal axis to the vertical. Other pairs of types only have cells expressing one type or the other, as in the L-shaped neutrophil-megakaryocyte histogram.

Our scTOP-based visualizations of developmental decisions during hematopoiesis complement the analysis in [46] using pseudo-time. Like in the original paper, we find strong evidence for the existence of progenitor cells that can give rise to multiple lineages (Figure 9), including a neutrophil/monocyte bivalent progenitor and possibly a basophil/eosinophil/neutrophils trivalent progenitor. The existence of these multi-potent progenitors seems to give rise to graded developmental dynamics where intermediate cells have significant projections on multiple cell types. This is in stark contrast with developmental decisions between other cell lineages (megakaryocytes vs. monocytes, basophils vs. megakaryocytes, monocytes vs. basophils) that seem to be mutually exclusive. One compelling feature of these visualizations using scTOP is that they require no statistical fitting, dimensional reduction, or ordering of cells. Once we have chosen the relevant reference basis (in this case, progenitors and six differentiated lineages), the resulting plots give an interpretable way of assessing complex developmental dynamics.

### C. Using scTOP to compare endogenous and transplanted cells

Recent advancements in cell culture and cell transplantation techniques have allowed researchers to generate cells that were maintained or differentiated in vitro to approximate and eventually replace endogenous cells generated during in vivo development. However, even with scRNA-seq, it is difficult to consistently quantify the transcriptomic similarity between donor-derived and endogenous lineages from transplant recipients. Since it provides a quantitative measure of cell type similarity, scTOP can be useful for comparing cell populations. A relevant example can be found in pulmonary cell replacement therapy, where one goal is to differentiate or maintain alveolar cells in vitro and then transplant them into injured lungs.

To compare two distinct pulmonary cell transplant protocols, we used scTOP to analyze the results of scRNA-seq from two recent publications featuring murine pulmonary cell transplantation [35, 47]. In Louie et al., adult mouse AT2 cells were maintained in culture with support cells for 3 weeks to generate a donor cell population. In contrast, in Herriges et al. the authors developed a protocol for directed differentiation of mouse PSCs into lung distal tip-like cells, which mimic an embryonic progenitor of AT2 cells. In both papers, the donor cells were transplanted into injured mouse lungs, where they survived and gave rise to multiple cell lineages, including AT2-like cells.

To highlight the usefulness of scTOP, we first consider another dimensional reduction method. Dimensional reduction methods like UMAP and t-SNE are often used to compare similar cells from different sources, such as transplanted and endogenous cells. Figure 12 shows the AT1 and AT2 cell populations from Herriges et al. and Louie et al., both donor-derived and endogenous. In figure 12 (a), three possible UMAP plots are shown with varying algorithmic parameters. The qualitative relations between the clusters are generally consistent, with AT1 cells clustering together apart from the AT2 cells. However, the distances between cells and the relative positions of clusters are not consistent when the UMAP parameter is varied. This illustrates how UMAP is useful for visualization and gaining qualitative intuition for a particular dataset but cannot be used to make a rigorous quantitative comparison between cells.

**FIG. 12.**
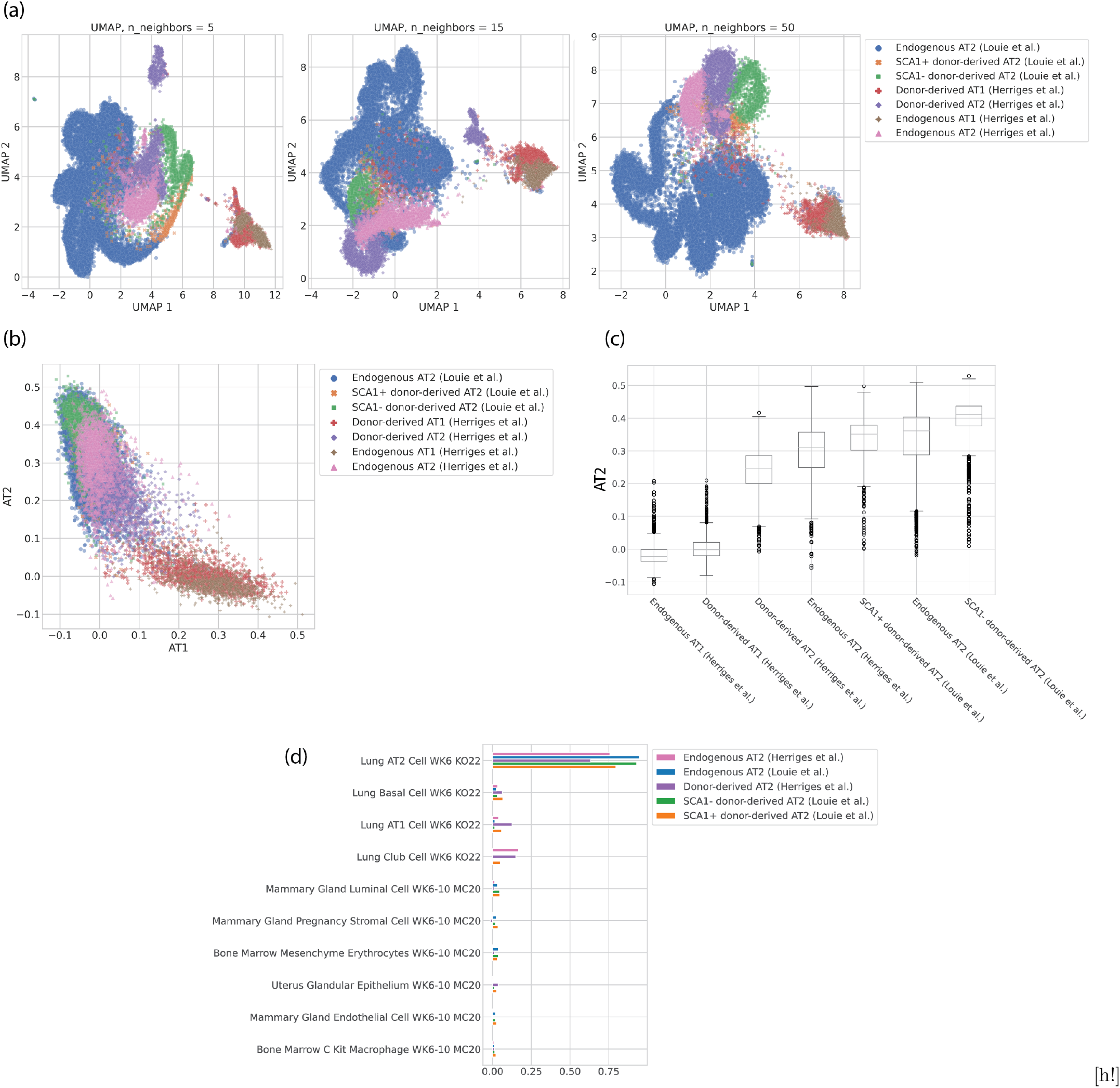
Donor-derived and endogenous alveolar cells from Herriges et al. and Louie et al. The cell type annotations come from a separate Louvain clustering and are the same for all plots in this figure. (a) UMAP visualizations of the data, colored by cell type annotations. The three have different values for the UMAP parameter n neighbors, which changes whether the embedding preserves more local or global structure. (b) scTOP plot showing the AT1 score on the x-axis and the AT2 score on the y-axis. (c) Box-and-whisker plot showing the distributions of scTOP AT2 scores for the various cell populations. (d) Bar plot showing the top ten aggregate scores for the AT2 populations from different sources, sorted by the scores for the SCA1+ population.

On the other hand, figure 12 (b) shows the scTOP scores for AT1 and AT2 for the same populations. The distances between cells will only change if the reference basis is changed, and even then will not change significantly (see S3 for discussion). The AT1 and AT2 populations are separated along the relevant axes, and the highly-similar cell populations are largely overlapping. Because scTOP provides many relevant dimensions to choose from instead of trying to capture high-dimensional distances in two dimensions, it is possible to directly compare different populations to the same cell type. Figure 12 (c) shows the distributions of AT2 scores for each of the populations, providing a clear quantitative way to compare the various engineered cells. As expected, the endogenous populations from both data sources are similar in distribution. The means and overall distributions of the AT2-derived (Louie et al.) cells are higher than those for the PSC-derived (Herriges et al.) cells, suggesting that the primary cell transplantation better emulates the endogenous AT2 cells.

Figure 12 (d) provides a comparison between cell populations using aggregate scTOP scores. Similar to figure 12 (c), (d) provides a quantitative comparison between the populations. This figure confirms that while both AT2-derived and PSC-derived transplants generate cells that have high AT2 scores, the AT2-derived cells are quantitatively more similar to the reference AT2 profile. Altogether this analysis demonstrates how scTOP can be used to compare two distinct cell populations against a primary control. scTOP satisfies the need for quantitative assessment of transplantation protocols and can be used in combination with functional tests to identify protocols that most effectively replace endogenous cell lineages.

## IV. DISCUSSION

In this paper, we have introduced a new physics-inspired method, scTOP, for analyzing scRNA-seq data. scTOP projection scores provide clear and meaningful axes to visualize differentiation and classify cells. Since scTOP requires no statistical fitting, clustering, or dimensional reduction techniques, plots can be made quickly and easily, even for an individual cell. The input to scTOP is a reference basis of cell fates of interest, allowing us to take advantage of the wealth of new scRNA-seq expression atlases now being generated. Importantly, scTOP scores are robust to the choice of basis and reproducible across experiments, labs, and datasets, allowing integration of disparate data sources. scTOP is especially suited for tasks where the cell types of interests are well characterized. For example, we have found in our own work that scTOP is an extremely sensitive measure for assessing the fidelity of engineered cells from directed differentiation protocols and reprogramming [35]. Since scTOP does not require data harmonization, joint embeddings, or make use of cellular neighborhood information, scTOP is able to distinguish between technical noise and biologically meaningful differences. scTOP also can be used to classify cells with minimal computational costs and thus represents a potentially simple, scalable, and reproducible way of dealing with the proliferation of scRNA-seq datasets. Since scTOP yields an alternative coordinate system for representing developmental dynamics in cell-fate space, scTOP-based visualizations are a powerful way of thinking about complex developmental processes.

As an illustration, we analyzed the development of the lung epithelial AT1 and AT2 cell fates [40] and developmental dynamics during hematopoiesis [46]. Using scTOP, we showed that it was easy to identify a transient hybrid AT2/AT1 population 3 to 5 days post-birth whose gene expression profile is distinct from previously investigated transient AT2-to-AT1 states. By plotting scTOP scores as a function of time for AT1 and AT2, hybrid AT2/AT1 cells were easily identified since they formed a distinct cluster that was well separated from mature AT1 and AT2 cells. This example illustrates how given an accurate reference basis, scTOP is able to pick up on small but biologically meaningful differences in global gene expression profiles.

In the context of hematopoietic development, where HSCs give rise to multiple downstream cell fates, we took advantage of the fact that scTOP maps each cell to a seven-dimensional coordinate (each direction in the coordinate systems corresponds to one of the seven cell types in the reference basis: undifferentiated progenitors, neutrophils, monocytes, megakaryocytes, mast cells, eosinophils, and basophils) to generate multiple two-dimensional visualizations using scTOP. We found that a single clonal family can often give rise to up to three different mature cell fates. Our visualizations also suggest that developmental decisions between different cell fates can be classified into two broad categories: mutually exclusive and graded. For cell fates such as megakaryocytes and monocytes, cells never have significant projection scores on both lineages simultaneously, suggesting that the underlying developmental decisions are mutually exclusive. In contrast, for other pairs of cell fates, such as monocytes and neutrophils, cells often exhibit significant projections on both members of the pair suggesting these developmental switches function in a more graded manner. It will be interesting to see if this general distinction is also present in other developmental systems or is specific to hematopoiesis.

In the analysis here, we have limited ourselves to considering scRNA-seq data. However, in principle, our approach can be easily extended to include other data modalities, such as chromatic accessibility and histone modifications. Recently, several interesting works such as Dynamo have used RNA-velocity to directly learn cellular dynamics [20, 48]. It will be interesting to see if these methods can be combined with scTOP to learn vector flows directly in cell-type space. Similarly, methods based on optimal transport, such as Waddington-OT [10], or developmental directory reconstruction based on graph embedding, such as Monocle [49], can also be performed in the cellular identity space instead of gene expression space. These represent promising directions for future research since the scTOP cell identity coordinates are often better suited for analyzing developmental dynamics than gene expression coordinates.

We have found that scTOP can be used to quantitatively assess the fidelity of cultured cells, as in the case of donor-derived lung alveolar cells. However, we currently have no systematic manner within the scTOP framework for providing guidance on adjusting gene expression or designing differentiation protocols to steer cells toward desired fates. Developing such methods would provide powerful new computational tools for engineering cells via directed differentiation.

We are also interested in extending scTOP to better understand the signaling pathways and genetic signatures underlying developmental dynamics. While scTOP provides a way to translate between gene expression space and cell type space, there is currently no direct interface with the biologically-essential signaling space. Further work in uniting the three relevant spaces, combined with studying the possible bifurcations in these spaces, may provide insight into the complex process of differentiation.

## ACKNOWLEDGMENTS

We acknowledge useful discussions with Jason Rocks, Robert Marsland, and Alex Lang. We would like to thank the Mehta group and Kotton lab for their comments on the manuscript. The work was funded by a grant to the Boston University Kilachand Multicellular Design Program (to DK and PM) and NIH NIGMS 1R35GM119461 to PM. We also acknowledge programming support from Alena Yampolskaya and Sumner Hearth.

## APPENDIX

### A. Preprocessing

The raw scRNA-seq count data for each dataset was downloaded as a matrix of mRNA counts corresponding to genes and cells. An example of such a matrix is shown in step 0 of figure S1, where each row is a different gene, and each column is a different cell. Throughout this figure, the histogram to the right of each step shows the distribution of entry values for the first column (cell) in the matrix, and the histogram below the step shows the distribution of entry values for the second row (gene Rpl7) of the matrix. The histograms are presented as examples of how the distributions (for each cell and gene) of gene expressions change as the data is preprocessed.

In the first step of preprocessing, each cell is normalized independently. This step is taken because scRNA-seq generally measures relative rather than absolute expression levels within a cell. In order to facilitate the comparison of cells across sequencing conditions, we transform the normalized RNA counts into quantities that are centered around the mean expression level. Z-scores are exactly such a quantity; they measure the number of standard deviations away from the mean. To calculate the z-scores after step 1, we convert the normalized gene expressions into percentiles by ranking each gene in a given cell from lowest to highest, then dividing by the number of genes (plus one so that the highest rank is not assigned the 100th percentile). For example, if a cell has 1000 genes sampled and Rpl7 is the highest-expressing gene, it will be given rank 1000 and labeled as the 99th percentile. If two or more genes are tied for rank, as is the case for zero-values, they are assigned the average of the ranks they would have been assigned if they were not tied (see rankdata function from the Python package Scipy). To convert from percentiles to z-scores, we assumed a normal distribution with mean 0 and standard deviation 1 and applied the corresponding quantile function.

### B. Reference basis construction

Creating an appropriate reference basis is vital to the accuracy of scTOP. There are many existing scRNA-seq atlases, such as the Mouse Cell Atlas; using them with scTOP requires some amount of curation. For each reference dataset, we took an average over each population corresponding to a given cell type. The average gene expression profiles were preprocessed and then used as the reference profiles for each cell type. Often, it was necessary to perform additional curation of the reference basis. Cell types were dropped from an atlas if they did not include enough cells (i.e., if they contained *<* 100 cells) or were found to be poorly defined (i.e., not enough marker genes to distinguish that cell type from others). Common instances of the latter were cell types that differed only in the expression of a single gene.

**FIG. S1.**
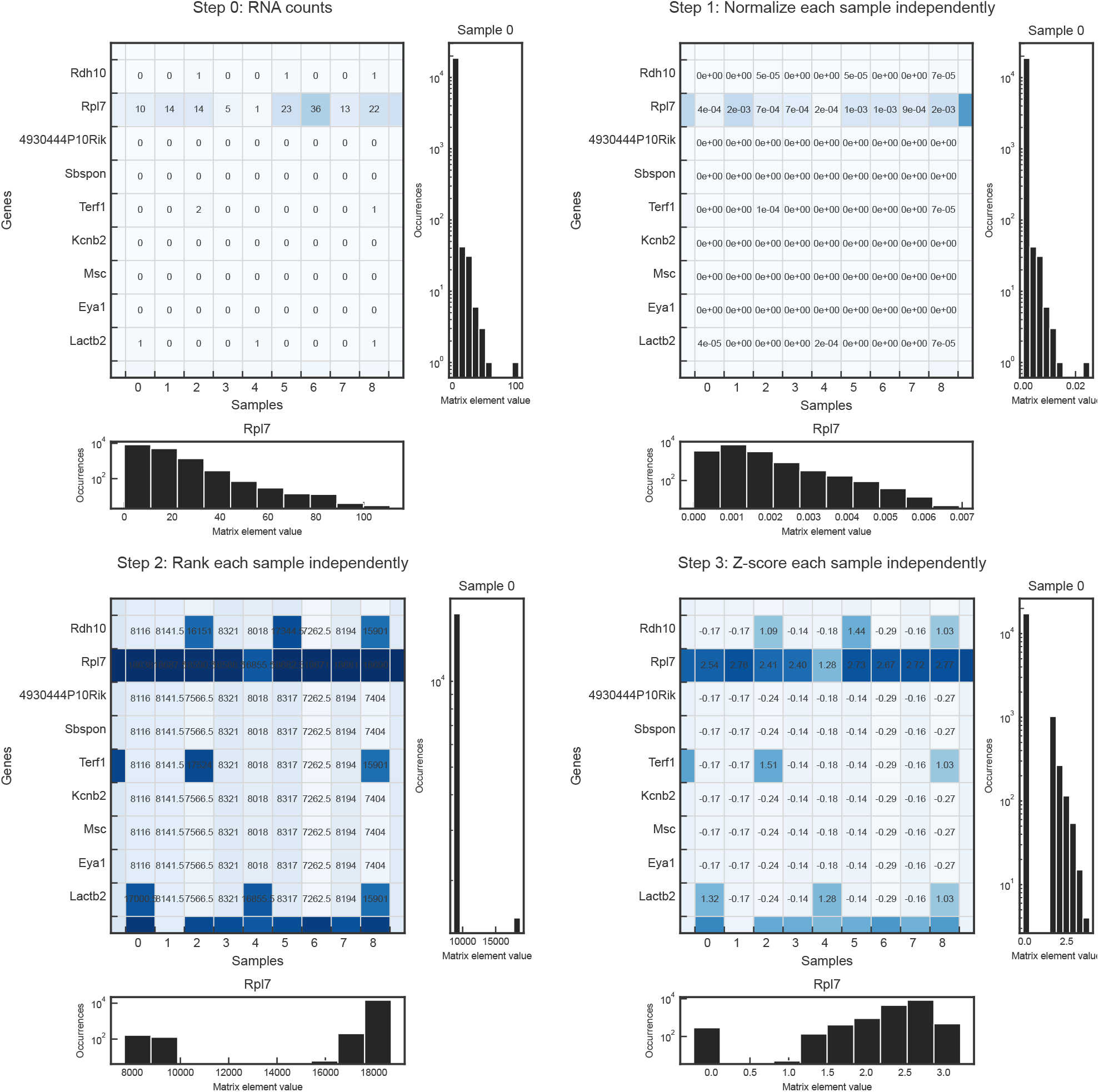
Data distributions at each preprocessing step. Step 0 shows the original RNA count matrix before preprocessing, where each entry corresponds to the number of RNA detected for a particular gene in a particular cell. The rows of each matrix correspond to genes, and the columns correspond to cells. The histograms to the right of each matrix show the distribution of an example cell, while the histograms below each matrix show the distribution of an example gene. Step 1 of preprocessing is to normalize each cell. Steps 2 and 3 convert the normalized gene expressions to z-scores. Step 2 ranks the genes in each cell according to magnitude and converts the rank into a percentile. Step 3 uses a quantile function, assuming a normal distribution with mean zero and standard deviation one, to convert from percentiles to z-scores.

Ensuring that each reference gene expression profile sampled enough cells was essential. This is because the number of cells determines how well-sampled the reference cell type is. Although cell types have distinct transcriptomic patterns, they are not all transcriptomically identical. There are variations in gene expression levels between cells of the same type due to differences in cell-level processes such as cell division or environmental stimuli. Averaging over a population of cells which are some cell type X creates an approximate gene expression profile of the archetypal X cell.

#### 1. Mouse Cell Atlas

The Mouse Cell Atlas (MCA) contains hundreds of thousands of cells from tissues across the mouse body at various stages of development. For the reference basis created from the Mouse Cell Atlas (MCA), we dropped cell types that had fewer than 100 cells. For cell types that had greater than or equal to 100 cells, we averaged across the entire population for each one, then preprocessed. For cell types that were divided according to high expressions of various genes, we combined them into one cell type. For example, the original MCA contained dozens of clusters of mammary gland secretory alveoli cells such as “secretory alveoli cell, Hes1 high” or “secretory alveoli cell, Gpx3 high.” These were all combined into one cell type, “mammary gland secretory alveoli.” This is because we wanted each reference basis type to act as an archetype rather than a perfect representation of every possible iteration of a cell type; each reference type represents a basin of attraction, and the basins were defined by the overall cell type, not whether individual genes were particularly high within that cell type. We cleaned each of the MCA type labels and ended up with cell types defined by the organ of origin, specific cell type, and time point of collection (e.g., “lung ciliated cell week 6-10”). The time points of cell types in the MCA were either E14.5 or somewhere between week 6 and week 10 post-birth. The resulting reference basis contained 221 cell types.

### C. Accuracy measures

For table I, the top1 accuracy was determined by applying scTOP to a cell, finding the type with the highest score, and checking whether that type matched the cell’s true type. Top3 was similarly determined, except instead of just checking the highest-scoring type, we checked whether the true type was included in the top 3 highest-scoring types.

By looking at the scTOP scores for cells whose true type did not match the reference type, we set a score of 0.1 as the lower bound for identifying a type. As shown in figure S3 (c) and (f), when the reference type does not match the sample type, the scores for individual cells fall well within [-0.1, 0.1], even when most of the reference basis is missing.

In table I, we labeled cells as unspecified if the highest scTOP score was less than 0.1. This provides a useful metric for determining whether a query cell belongs to one of the cell types in the reference basis. If the highest scTOP score is lower than 0.1 for a query cell, this indicates that the reference basis lacks the relevant types or was poorly constructed (see appendix IV D 1 for a discussion of ill-defined bases).

scTOP can only identify cells that exist within the reference basis. As such, the accuracy scores were calculated only for the cells whose true type (as determined by the annotations from the authors) was included in the reference basis for the corresponding analysis. For cells whose true types were not represented in the basis, the unspecified rate was very high. This indicates that the rate of false positives is very low.

### D. Robustness of scTOP

scTOP is consistent between iterations of the algorithm, and it is robust to changes in the reference basis. As shown in figure S3, many scTOP scores do not change significantly even when most of the reference basis is removed. Generally, scores for cells that match the reference type do not change much even when only 25% of the original basis is retained (figure S3 (d), (e)). The largest difference occurs for cells that are similar to the reference type, such as the AT1 score for AT2 cells (figure S3 (g)). These scores tend to increase because removing other types from the basis reduces the ability to de-correlate effectively. In other words, scTOP is more likely to confuse similar types when it has fewer reference types to compare. This effect is most apparent when relevant types are removed from the basis. Figures S3 (j) and (k) show that all lung types score higher for AT1 and AT2 when only lung alveolar types are included in the basis. Figures S3 (c) and (f) show that irrelevant scores, such as bladder and thymus cells, do not change much even when large portions of the basis are removed. scTOP does not confuse sample cells for types that are dissimilar even with a small reference basis because there is no need to de-correlate between types that are naturally not correlated.

#### 1. Cases where it does not work

The performance of scTOP relies on the quality of the reference basis. scTOP is expected to give inaccurate results in cases where the reference basis is ill-defined. The underlying assumption of the algorithm is that the reference basis consists of accurate gene expression profiles of cell types that are attractor basins. In other words, the reference types are assumed to represent stable cell types rather than cell states that are on the path to the true cell type basins. For example, scTOP performs poorly when using a reference basis consisting of early embryonic identities because the cell types are not sufficiently differentiated from one another. Adapting scTOP to such early lineage data is an important and interesting problem.

The other part of the assumption is that the reference types should be accurate profiles of the cell type of interest. As discussed in appendix IV B, reference types are created from averaging over cells, and the more cells included in this average, the better-sampled the type is. As expected, scTOP gives unreliable results when the reference gene expression levels do not accurately reflect the true cell types’ gene expression levels. An important factor in the accuracy of scRNA-seq is the amplification step: after RNA is extracted, it is amplified to improve the sensitivity of sequencing, but not all RNA is amplified at the same rate. This results in inflated counts for some genes but not others. Unique molecular identifiers (UMIs) are vital for reducing the inflating effect of amplification [50]. UMIs tag the RNA before amplification and are present post-amplification; this makes it possible to see which amplified RNA came from the same original molecule and prevents uneven double-counting of RNA. We have found that scTOP gives more accurate results when used with reference and query datasets that use UMIs, compared to scTOP results using data that did not use UMIs.

### E. AT1/AT2 marker genes

In figure 7, plots in (b) and (c) were colored according to AT1 and AT2 marker gene expression. For each cell, the average was taken over the expressions of a set of marker genes. The AT1 marker gene set was:

- Ager
- Akap5
- Aqp5
- Cldn18
- Clic3
- Clic5
- Col4a3
- Col4a4
- Cryab
- Cyp2b10
- Emp2
- Fam189a2
- Gprc5a
- Hopx
- Hs2st1
- Igfbp2
- Krt7
- Lmo7
- Mal2
- Pdpn
- Prdx6

**FIG. S2.**
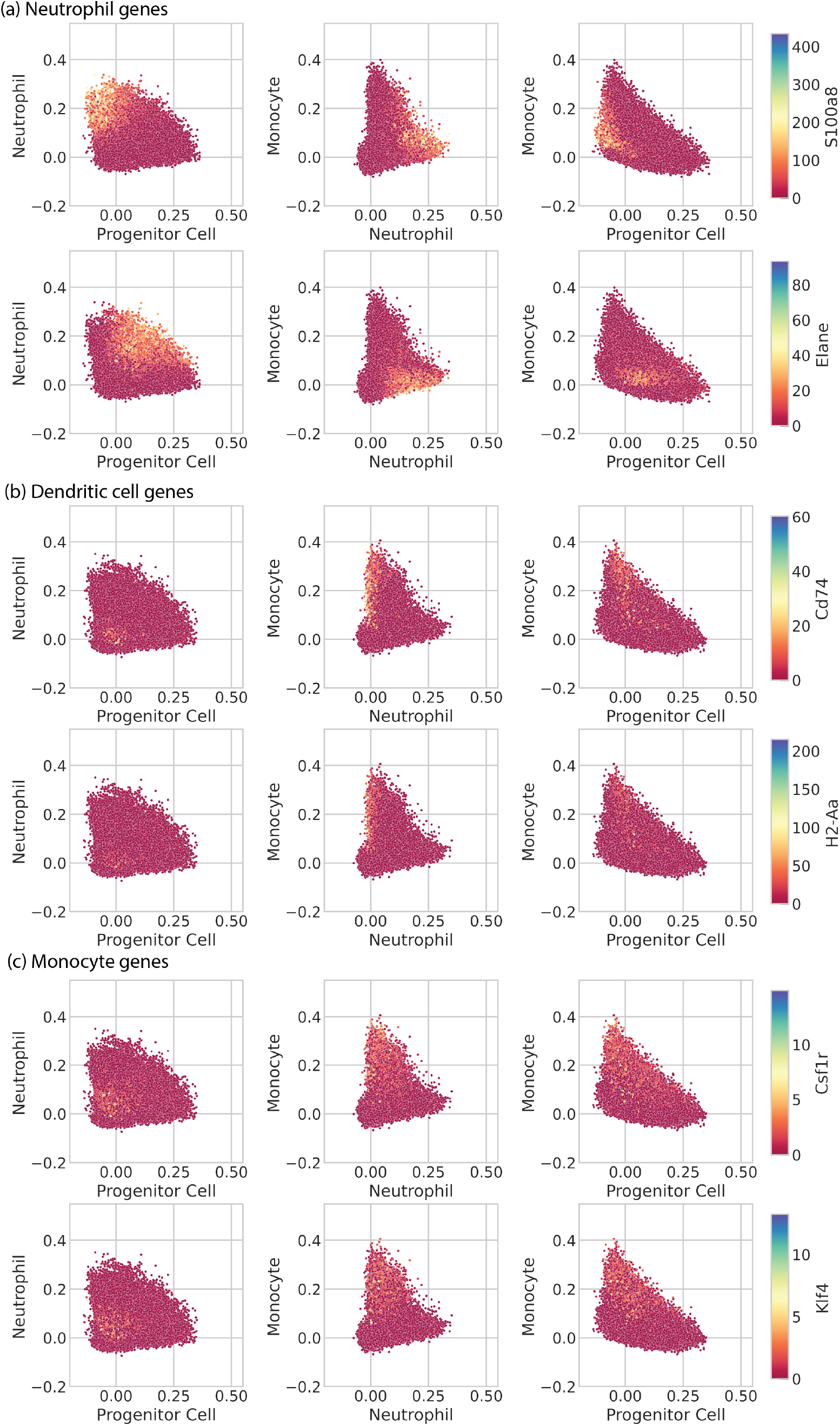
Scatter plots of in vitro hematopoietic clone families from Weinreb et al., with scTOP scores for progenitor, neutrophil, and monocyte types, and colored by normalized gene expression. In Weinreb et al. figure 4 (a), cells visualized using pseudo-time inference were separated into branches. The monocyte branch appeared to have half of its cells expressing dendritic cell genes and the other half expressing neutrophil genes. This figure plots those same cells on progenitor, neutrophil, and monocyte axes. The colors correspond to neutrophil, dendritic cell, and monocyte marker genes. (a) Hematopoietic cells colored according to the expression of neutrophil marker genes S100a8 and Elane. (b) Hematopoietic cells colored according to the expression of dendritic cell marker genes Cd74 and H2-Aa. (c) Hematopoietic cells colored according to the expression of monocyte marker genes Csf1r and Klf4.

- Pxdc1
- Rtkn2
- Scnn1g
- Spock2
- Tspan8
- Vegfa

The AT2 marker gene set was:

- Abca3
- Acot7
- Acoxl
- Acsl4
- Ank3
- Atp8a1
- Cd74
- Cebpa
- Cpm
- Ctsh
- Cxcl15
- Dram1
- Egfl6
- Elovl1
- Etv5
- Fabp5
- Fasn
- Hc
- Lamp3
- Lgi3
- Lpcat1
- Lyz2
- Mlc1
- Muc1
- Napsa
- Npc2
- Ppp1r14c
- S100g
- Scd1
- Sfta2
- Sftpa1
- Sftpb
- Sftpc
- Sftpd
- Slc34a2
- Zdhhc3

---

**FIG. S3.**
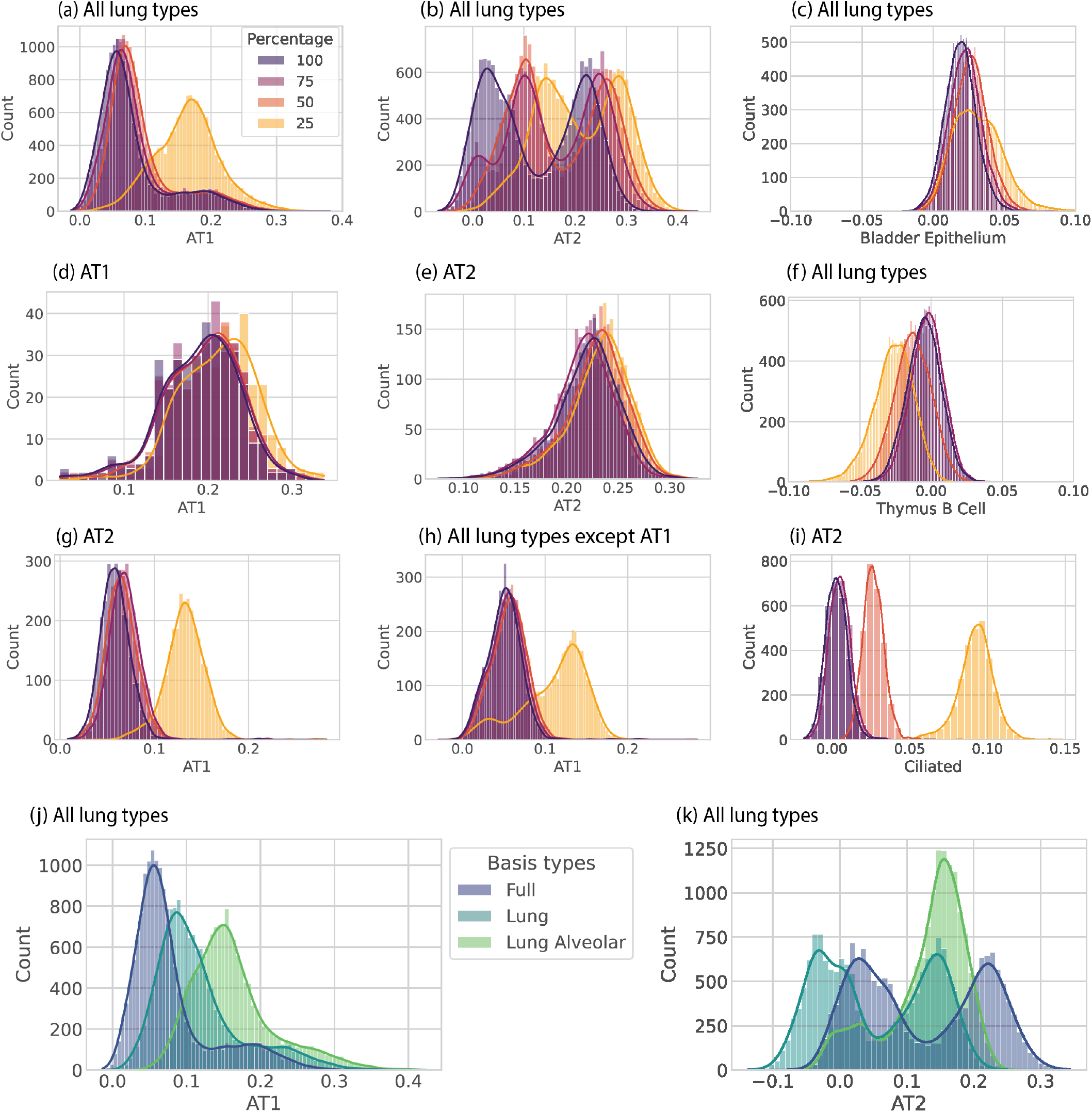
Distributions of scTOP scores for lung cells from Herriges et al., using varying portions of the Mouse Cell Atlas as the basis. The label above each plot indicates the Herriges et al. sample subset, while the x-axis label indicates the scTOP reference type. “All lung types” means that all of the lung types from the data set were included. (a) - (i) show how scTOP scores change when 25-100% of the basis is used. (j) - (k) show how scTOP scores change when the basis is changed from the full basis (which includes many tissues and organs) to just the lung types and to just the two alveolar types. (c) and (f) show scTOP scores for random irrelevant types, Bladder Epithelium and Thymus B Cell, to show that the scTOP scores remain low for types that do not match the sample.

